# Predominance of *cis*-regulatory changes in parallel expression divergence of sticklebacks

**DOI:** 10.1101/412932

**Authors:** Jukka-Pekka Verta, Felicity C. Jones

**Affiliations:** Friedrich Miescher Laboratory of the Max Planck Society, Max-Planck-Ring 9, 72076 Tübingen, Germany; Organismal and Evolutionary Biology, University of Helsinki, Viikinkaari 9, 00790, Helsinki, Finland

**Keywords:** gene regulation, parallel evolution, adaptation, selective sweeps, allele specific expression, genetic assimilation, additivity, mode of inheritance, Haldane’s sieve, epistasis

## Abstract

Regulation of gene expression is thought to play a major role in adaptation but the relative importance of *cis*- and *trans*-regulatory mechanisms in the early stages of adaptive divergence is unclear. Using RNAseq of threespine stickleback fish gill tissue from four independent marine-freshwater ecotype pairs and their F1 hybrids, we show that *cis*-acting (allele-specific) regulation consistently predominates gene expression divergence. Genes showing parallel marine-freshwater expression divergence are found near to adaptive genomic regions, show signatures of natural selection around their transcription start sites and are enriched for *cis-*regulatory control. For genes with parallel increased expression among freshwater fish, the quantitative degree of *cis-* and *trans-*regulation is also highly correlated across populations, suggesting a shared genetic basis. Compared to other forms of regulation, *cis-*regulation tends to show greater additivity and stability across different genetic and environmental contexts, making it a fertile substrate for the early stages of adaptive evolution.

## Introduction

The ability of organisms to rapidly adapt to new environments can be both facilitated and constrained by the underlying molecular basis and mechanisms operating at the genomic level. There have been significant advances in our understanding of the genomic basis of adaptive evolution including that adaptation is often polygenic and involves loci that are predominantly intergenic and putatively regulatory (Brawand et al., 2014; Grossman et al., 2013; Jones et al., 2012). Transcriptional regulation of gene expression can be controlled through *cis*-acting regulatory elements that are linked to their target gene alleles (e.g. promotors, enhancers), or through *trans*-acting mechanisms such as transcription factors whose action typically impacts both target alleles.

During rapid adaptation, selection may favour master regulator genes that set off concerted changes in many downstream genes within a gene regulatory network through *trans-*acting mechanisms. This view has theoretical backing and has been demonstrated in multiple examples (Cooper, Rozen, & Lenski, 2003, Stern & Orgogozo, 2009). Alternatively, selection may favor the modularity and tighter linkage offered by *cis*-acting mechanisms under the following scenarios: moderate changes to single alleles in a tissue-specific manner, rather than systemic changes to gene expression; or frequent out-crossing or hybridisation (Carroll, 2008). *Cis*- and *trans*-regulatory mechanisms are not mutually exclusive and adaptation is expected to promote co-evolution between *cis*- and *trans*-acting mechanisms so that optimal gene expression levels are reached and maintained (Fraser, Moses, & Schadt, 2010). Interdependence of *cis*- and *trans*-regulatory mechanisms has been hypothesized to act as a barrier for gene flow and contribute to incipient speciation: incompatible regulatory factors fail to promote optimal gene expression levels in hybrid progeny (Landry, Hartl, & Ranz, 2007a; Landry, Wittkopp, Taubes, Ranz, Clark, & Hartl, 2005).

Given their different properties, it stands to reason that different evolutionary scenarios and selection contexts may alternatively favour *trans-* and *cis-*acting mechanisms in the early stages of intraspecific adaptive divergence (Coolon, McManus, Stevenson, Graveley, & Wittkopp, 2014; Fraser et al., 2010; Hart, Ellis, Eisen, & Miller, 2018; Lemos, Araripe, Fontanillas, & Hartl, 2008; Stern & Orgogozo, 2009). In this context, divergent adaptation to local environments often occurs in the face of ongoing gene flow. The shuffling effects of recombination will tend to dissociate co-evolved factors and selection may be more efficient on the larger-effect haplotypes carrying multiple such factors that are maintained by linkage-disequilibrium. Thus the advantage of a rapid adaptive response mediated via a small number of *trans*-regulatory mutations in a gene regulatory network, may shift to favor *cis*-regulatory architecture where co-evolved mutations are more closely linked to each other and the gene whose expression they regulate. Parallel evolution provides a powerful context to explore the relative importance of *cis*- and *trans*-regulation in the early stages of intraspecific adaptive divergence. Using independent biological replicates of the evolutionary process it is possible to ask whether the same phenotype has evolved via the same or different molecular underpinnings. While regulatory changes seem to predominate in adaptation of natural populations we know little about the extent and parallelism in gene expression and its *cis*- and *trans*-regulation.

The threespine stickleback fish is an excellent system to address these questions. Following the retreat of the Pleistocene ice sheet 10-20k years ago the parallel evolution of freshwater ecotypes from ancestral marine forms has occurred repeatedly and independently in thousands of populations across the Northern Hemisphere (Bell & Foster, 1994). Considerable evidence points towards an important role for gene regulation in this adaptive divergence. Firstly, forward mapping and functional dissection have identified mutations in *cis*-regulatory elements underlying the parallel loss of major morphological traits (bony armor plates Colosimo et al., 2005; O’Brown, Summers, Jones, Brady, & Kingsley, 2015 and pelvic spines Shapiro et al., 2004; Chan et al., 2010). Further, whole genome sequencing of marine and freshwater sticklebacks from multiple populations revealed that repeated parallel evolution of freshwater ecotypes from marine ancestors involves reuse of pre-existing genetic variation at ~81 loci across the genome that are repeatedly involved in parallel evolution (Jones et al., 2012). These loci are predominantly intergenic and thus may act through regulatory mechanisms.

During adaptation to their divergent environments marine and freshwater sticklebacks have evolved differences in numerous morphological, physiological and behavioral traits. Two key divergent traits include their anadromous (migratory marine) versus resident-freshwater life histories and the ability to live in fresh- and saltwater. For this adaptation, the gill’s role in osmoregulation and respiration is likely to be particularly important (e.g. through regulation of ion channel genes, Evans, Piermarini, & Choe, 2005). In saline water, fish counteract water loss and ion gain by ion exclusion. In freshwater, fish compensate against ion loss and water gain by ion uptake. Expression changes in osmoregulatory genes has been linked to freshwater adaptation by anadromous ancestors in sticklebacks and other fish (Gibbons, Metzger, Healy, & Schulte, 2017; Velotta et al., 2016). Further, previous studies have shown that freshwater adaptation in sticklebacks and other fish is associated with changes in gene expression plasticity (Gunter, Schneider, Karner, Sturmbauer, & Meyer, 2017; McCairns & Bernatchez, 2010; Whitehead, 2010). The genetic basis of these gene expression differences is not known but leads to the prediction that loci showing parallel divergence in gene expression levels show parallel and environmentally insensitive mechanisms of gene regulation in independently evolved marine and freshwater ecotypes.

Here we study the evolution of gene expression and its *cis*- and *trans*-regulation in the gills of threespine sticklebacks as a model for regulatory evolution during early stages of parallel adaptive divergence with gene flow. Using freshwater-resident and anadromous marine sticklebacks from rivers in Scotland (3) and Canada (1), we ask to what extent parallel divergent adaptation to marine and freshwater environments involves parallel expression divergence in the gills under standardized laboratory conditions. We explore whether parallel differentially expressed genes are found more frequently near previously identified adaptive loci and whether they show molecular signatures of natural selection. We then dissect the *cis*- and *trans*-regulatory basis of gene expression differences using allele specific expression analysis of marine-freshwater F1 hybrids and their parents. We ask whether *cis*- or *trans*-regulatory changes predominate in the early stages of adaptive divergence with gene flow, and, by comparisons across marine and freshwater ecotypes from four independently evolving river systems examine the degree of parallelism in *cis*- and *trans*-architecture. Finally, by rearing F1 siblings in different water salinity conditions we explore the extent to which the degree of *cis*- and *trans*-regulation of divergently expressed genes is influenced by the environment.

## Results

### Stickleback gill transcriptome assembly

We analysed gene expression in the gill of marine and freshwater ecotype pairs each collected and derived from four river systems in Scotland and Canada (Fig 1a, Table S1). Gills of mature and reproductively active first-generation wild-derived female and male fish were dissected and their transcriptomes analyzed using strand-specific RNA-seq. We built a reference-guided assembly (Trapnell et al., 2012) of the stickleback gill transcriptome based on RNA-seq reads from 10 freshwater and 10 marine fish from 4 marine and 4 freshwater strains (Table S2). The stickleback gill transcriptome contains 29295 transcribed loci, 17304 of which are multi-transcript loci with 171620 different transcripts combined. This is considerably more than Ensembl gene build 90 for the stickleback genome (22456 loci, 29245 transcripts), likely because of the higher sequencing coverage of the current dataset compared to the low coverage EST libraries used to inform gene annotations of Ensembl’s genebuild (Kingsley et al., 2004). The number of transcripts per locus is highly skewed with a median of 3 and a small number of loci, including genes with immune function, with very high numbers of splice forms (Fig S1). The transcriptome includes 7147 novel transcribed loci that do not overlap with any transcript in Ensembl gene build v90. Of these novel loci, candidate coding regions with complete open reading frames and likelihood scores >20 were identified for 1018 using TransDecoder (Haas et al., 2013). At the locus level our assembly shows very high sensitivity (Sn=81%) with few false negatives (fSn=100%) and moderate specificity reflective of the appreciable number of novel coding regions we detected relative to the Ensembl gene build 90 (Sp=59%, fSp=69%). Considering the raw FPKM data 21399 (73%) of loci were expressed at FPKM>=1 in at least one of the 20 marine and freshwater fish analyzed, and 16195 (55%) in more than ten individuals. Hierarchical clustering of expression levels revealed that the gill transcriptome can be characterized by five major groups of loci based on their average expression level (Fig S2) with the most highly expressed group of genes showing strong enrichment for biological processes with the respiratory function of the gill including mitochondrial respiration, ATP synthesis coupled proton transport and cytoplasmic translation (Supplementary note).

**Figure 1.**
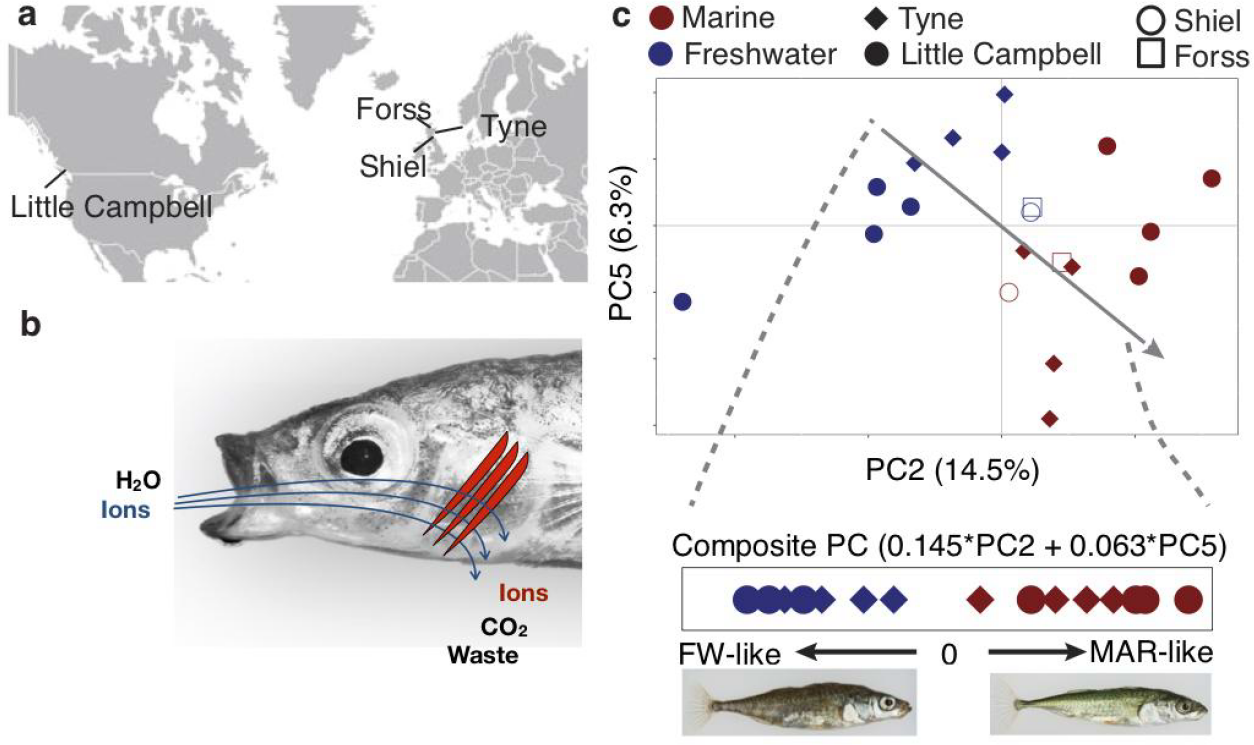
Freshwater and marine sticklebacks show parallel expression divergence among a largely non-parallel evolving transcriptome. (**a**) Marine and freshwater strains sampled from 4 different river systems (**b**) The gill is a multifunctional organ with roles in osmoregulation, respiration and waste excretion. In freshwater gills uptake ions (blue), while in saline water ions are excreted (red). (**c**) Principal components analysis of normalized expression levels separates marine (red) from freshwater (blue) ecotypes along a composite PC axis (grey line). PCA is calculated based on a sample-size balanced set of Tyne and Little Campbell samples (solid symbols), onto which Forss and Shiel individuals are projected (open symbols).

### Detecting parallelism in marine-freshwater transcriptome divergence in a largely non-parallel evolving transcriptome

We hypothesized that selection for divergent adaptation to marine and freshwater habitats could drive parallel divergence in gene expression in multiple independently evolving marine-freshwater ecotype pairs of distinct geographic origin. To test this hypothesis, we first investigated the major sources of covariation in the freshwater and marine transcriptomes using Principal Component Analysis (PCA).

While the first major axis of variation separates individuals by river system (24% variation explained Fig S3), we identified PC2 and PC5 as major axes of variation (14.5% and 6.3% PVE respectively, Fig 1c) that separate marine and freshwater transcriptomes. We defined a composite PC axis that captures parallel divergence in the transcriptomes of freshwater and marine ecotypes by summing the eigenvalue-weighted loadings from PC2 and PC5 (Fig 1c). The loadings of each transcript on this composite PC were used as a measure of each transcript’s individual contribution to parallel transcriptome divergence. Parallel transcriptome divergence is highly correlated with river-specific measures of marine-freshwater transcriptome divergence (log of marine/freshwater expression fold change in the River Tyne and Little Campbell Rivers, Fig S4). Similar to the only 0.01% parallel genetic divergence observed at the genomic DNA level (Jones et al., 2012), here parallel divergence of gene expression between marine and freshwater ecotypes reared under common environmental conditions represents only a small proportion of transcriptome variation. From here on we refer to loci with the highest contribution towards parallel marine-freshwater expression divergence (top or bottom 1% composite PC loadings, N=586 transcripts) as ‘*parallel diverged loci’*.

The parallel diverged loci show an enrichment in gene ontology processes and molecular functions associated with gill ion exchange, osmoregulation and blood traits (similar results were obtained from a differential expression analysis using Cufflinks, see Supplementary Note, Fig S5, Table S3). Among the overrepresented categories were multiple processes involved in ion transmembrane transport, suggesting that the parallel diverged transcripts function in regulating osmolarity through ion exchange. Top ranking genes include Na-Cl cotransporter (slc12a10), Basolateral Na-K-Cl Symporter (slc12a2), cation proton antiporter 3 (slc9a3.2), Potassium Inwardly-Rectifying Voltage-Gated Channel (kcnj1a.3), potassium voltage-gated channel (KCNA2), Epithelial Calcium Channel 2 (trpv6), Sodium/Potassium-Transporting ATPase (atp1a1.4), aquaporin 3a (aqp3a), and carbonic anhydrase 2 (ca2) — genes known to play a role in osmoregulation in fish and other organisms. We also note that our analyses identified differentially expressed novel loci that have no overlap with gene annotations from the Ensembl gene build. These results are consistent with our hypothesis that adaptive expression divergence influences physiological functions of the gill associated with a transition to permanent freshwater environment.

### Natural selection on parallel expression divergence

To explore the role of natural selection in parallel expression divergence, we used adaptive loci identified in a previously published study (Jones et al., 2012) and analysed newly generated whole genome sequence data of six unrelated fish of each ecotype from both the River Tyne and Little Campbell River for molecular signatures of selective sweeps.

Transcripts with parallel expression divergence are distributed across all 21 chromosomes (Fig 2a), and show significant proximity to genomic regions undergoing parallel marine-freshwater adaptive divergence at the DNA sequence level (identified in (Jones et al., 2012), randomization test, P<<0.025; Fig 2b-c). More than 13 percent of transcripts with parallel expression divergence were found within 10kb of regions of parallel genetic divergence; and nearly 40% within 75kb (Fig 2c), consistent with the hypothesis that the predominantly intergenic marine-freshwater adaptive loci identified in Jones et al., (2012) contain regulatory elements contributing to parallel marine-freshwater divergence in expression. We found similar results when we calculated parallel genetic divergence (CSS) based on the whole genome sequences generated in this study (Supplementary note, Fig S6).

**Figure 2.**
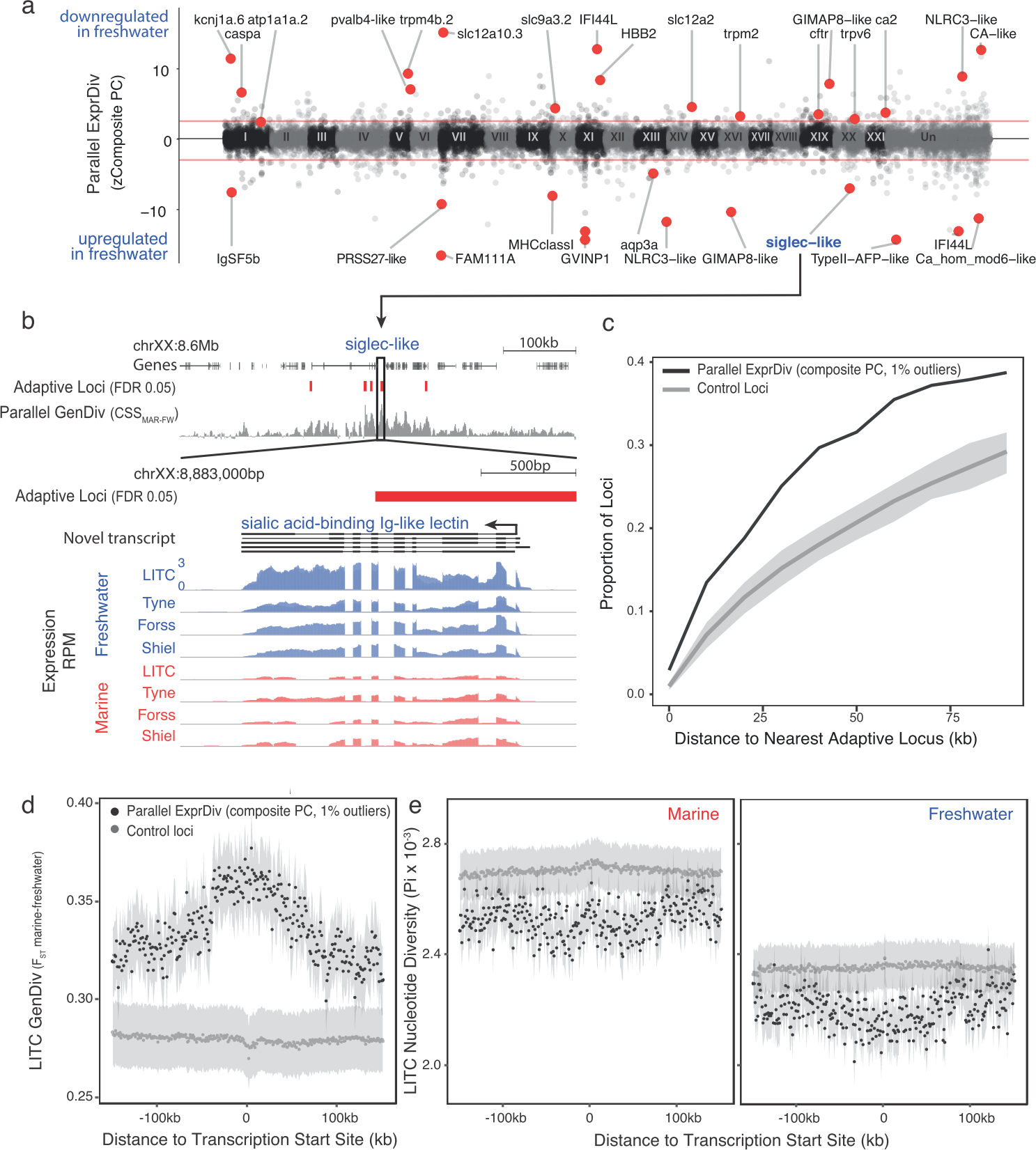
Genes with parallel marine-freshwater expression divergence are proximal to genomic regions underlying marine-freshwater adaptive divergence and show molecular signatures of natural selection. (**a**) Z-standardized composite PC loadings (zComposite PC) for each analyzed transcript across the genome highlighting transcripts with strong parallel marine-freshwater divergence in gene expression. Red lines correspond to the upper and lower 1% composite PC quantiles. Genes highlighted with red points have putative roles in ion transport and osmoregulation *(kcnj1a.6, atp1a1a.2, pvalb4, trpm4b.2, slc12a1, slc12a10.3, slc9a3.2, trpm2, trpv6, cftr, ca2, CA-like, aqp3a), calcium homeostasis (FAM111A, ca_hom_mod6-like), respiration (HBB2), cold temperature adaptation (TypeII AFP), jaw, gill and skeletal morhpogenesis (caspa, FAM111a), and immune system function (IFSF5b, caspa, PRSS27, MHCclassI, GVINP1, IFI44L (x2), GIMAP8-like (x2), NLRC3-like (x2), siglec).* (**b**) An example highlighting the proximity of parallel differentially expressed genes to adaptive loci is a novel transcript coding for a sialic-acid binding Ig-like lectin. This gene shows strong parallel expression divergence among freshwater and marine ecotypes from all four river-systems and overlaps a previously identified adaptive locus with a signature of parallel marine-freshwater genomic divergence (CSS FDR 0.05, (Jones et al., 2012)) (**c**) Across the genome, loci with strong parallel expression divergence (loci in the upper and lower 1% quantiles of composite PC, black line) are more proximal to adaptive loci identified in Jones *et al* 2012 than randomly sampled “control” loci (grey line). Grey shading shows 95% confidence intervals from 100 random samples of 586 transcripts. (**d & e**) In marine and freshwater fish from the Little Campbell River (LITC), loci with strong parallel expression divergence show molecular signatures of natural selection (d, increased genetic divergence FST; and e, decreased nucleotide diversity, Pi) centered around their transcription start sites (black points) compared to control loci (expressed loci showing non-parallel expression divergence). Points represent mean values of 10kb sliding windows and grey shading shows the standard error of the mean.

Natural selection is expected to reduce the diversity of a local genomic region leaving detectable molecular signatures of selection such as increased genetic divergence (FST) and reduced diversity (Pi) around adaptive loci (Nielsen et al., 2005). We calculated genetic divergence and nucleotide diversity flanking transcription start sites (TSSs). In Little Campbell fish we observed a reduction in nucleotide diversity (Pi) around the TSSs of transcripts with parallel expression divergence (Fig 2f-g). Reduced within-population diversity was accompanied by increased genetic divergence (FST) that was centered on TSSs with a slight upstream bias (Fig 2e). In contrast, although we observed a slightly increased FST around the TSSs of transcripts with parallel expression divergence in the River Tyne, we did not detect a reduction in nucleotide diversity relative to other transcripts (Fig S7), possibly due to concurrent selective sweeps on different haplotype backgrounds (also referred to as soft sweeps) in this younger population. Analysis of parallel differentially expressed transcripts defined using parametric tests gives similar results (Supplementary Note, Fig S8).

Combined, these results provide evidence for natural selection acting on parallel differentially expressed transcripts in the gill. This confirms the gill organ to be an important target of selection in the divergent adaptation of marine and freshwater sticklebacks to their respective environments and is consistent with gene regulation contributing to increasing genome-wide adaptive divergence of marine and freshwater sticklebacks.

### Expression divergence between stickleback ecotypes is predominantly *cis-*regulated

We hypothesized that the observed molecular signatures of selection around genes with divergent expression results from natural selection acting on *cis*-regulatory elements controlling gene expression levels. To investigate the role of *cis*-regulation, we compared the level of gene expression divergence in the gill transcriptomes of marine and freshwater parents to the level of allele-specific expression (ASE) in their reproductively mature F1 hybrid offspring using laboratory reared freshwater and marine strains from four independent river systems (Tyne, Forss, Shiel and Little Campbell).

Since F1 hybrids carry a marine and freshwater copy of each chromosome within a shared *trans* environment in each cell, any marine or freshwater allele-specific bias in transcript expression can be attributed to *cis*-acting regulatory elements, rather than *trans*-acting factors. We sequenced the parental genomes and identified coding-regions containing polymorphisms that were fully informative for allele-specific dissection (where the parents of a given cross were homozygous for alternate alleles, Fig S9, Fig S10). We classified regulatory divergence into *cis*-acting (allele-specific), *trans*-acting (non allele-specific) categories, or a combination of the two, based on the allele-specificity and magnitude of expression divergence in F1 hybrids compared to marine and freshwater parents (Fig 3a-c, Table S4) (Landry, Wittkopp, Taubes, Ranz, Clark, & Hartl, 2005a; Wittkopp, Haerum, & Clark, 2004). Since we are interested in genetic variation in natural populations, we used first generation wild derived parents and analyzed four F1 offspring per parental pair, in order to account for the different genetic backgrounds of each offspring. Considering all expressed loci in the gill, 3807 (13%), 4472 (15%), 7716 (26%) and 10102 (34.5%) non-sex-biased genes could be assayed for *cis*-vs *trans*-mediated expression in at least one F1 individual from Tyne, Shiel, Forss and Little Campbell, respectively.

**Figure 3.**
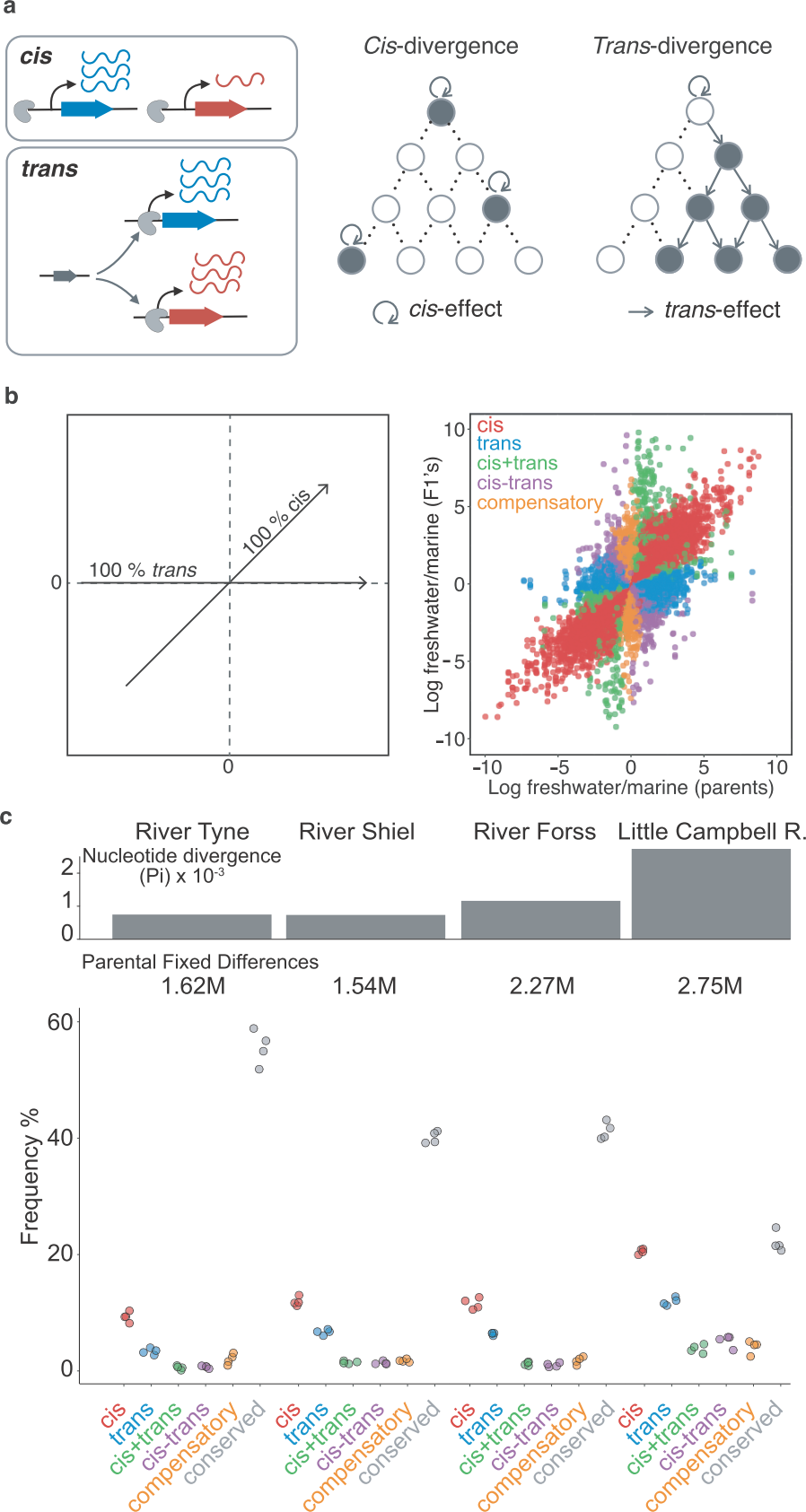
Expression divergence between marine-freshwater ecotype pairs is predominantly *cis-*regulated. (**a**) In F1 hybrids, allele expression is measured in the same *trans*-acting background, allowing dissection of expression variation due to effects that co-segregate with the allele (such as enhancers and promoters) from effects that influence both alleles and can thus be attributed to *trans*-acting sources (such as transcription factors). *Cis-* and *trans-*acting divergence is represented as effects that are contained within the nodes of a gene regulatory network (*cis*) or diffused from upstream regulators (*trans*). (**b**) Regulatory divergence is categorized in six categories by comparison of the expression log_2_-ratio of the marine and freshwater parent to marine- and freshwater-allele-specific expression log_2_-ratio within an the F1 offspring (Landry, Wittkopp, Taubes, Ranz, Clark, & Hartl, 2005). Individual data-points correspond to allele-specific expression values (y-axis) for each gene in each of four F1s relative to their parents (x-axis). Genes are colored according to their classification into different genetic architectures of expression divergence (red *cis*, blue *trans*, green *cis+trans*, purple *cis-trans*, gold *compensatory*, grey *conserved*. “*Ambiguous*” and “*conserved*” classes are omitted for clarity.) (**c**) *cis-*regulation is the predominant regulatory mechanism underlying gene expression divergence in all four river systems. Points show the overall frequency of regulatory divergence types, relative to the number of analyzed transcripts for each F1 offspring analysed. The proportion of *cis-*regulated expression divergence scales positively both with nucleotide divergence (grey bars) and the number of fixed differences (numbers beneath bars) between the marine and freshwater parents. “*Ambiguous*” class is omitted for clarity.

In each ecotype-pair, marine-freshwater expression differences are predominantly *cis*-regulated (Fig 3b, Fig S11), with the degree of *cis*-regulated expression divergence scaling positively with the degree of genetic divergence (measured either as the number of fixed differences or mean per site nucleotide divergence between the parents) between marine and freshwater ecotypes from each river system. Across the four river systems, from ~350 to ~2000 transcripts were classified as *cis-*regulated (with Little Campbell River fish showing the highest number, ~2000, or ~20%, of *cis-*regulated expression divergence. See also Fig. S12).

### Loci with parallel expression divergence are enriched for *cis*-regulation

We next tested whether particular types of gene expression regulation were more likely to contribute to marine-freshwater expression divergence that has evolved in parallel in multiple river systems. We used “parallel diverged loci” defined by the composite principal component described above that also show marine-freshwater expression differences in the same direction between the parents of our crosses and contain fully informative SNPs to allow for allele-specific analysis. After filtering, 181 and 68 parallel diverged loci were testable in Little Campbell and Tyne, respectively. Loci with parallel divergence in expression showed a significant excess of *cis-*regulation (18-29% and 20-22% above average in Tyne and Little Campbell F1 offspring respectively) and *cis+trans-*regulation (above average), compared to random expectations obtained by 1000 random draws of equal size (528 genes) from all genes analysed for allele specific expression (Fig 4 and Fig S13). In contrast, the overrepresentation of *trans*-acting divergence was lower, between 8-18% and 6-10%, respectively, and not different from random expectation in all F1’s.

**Figure 4.**
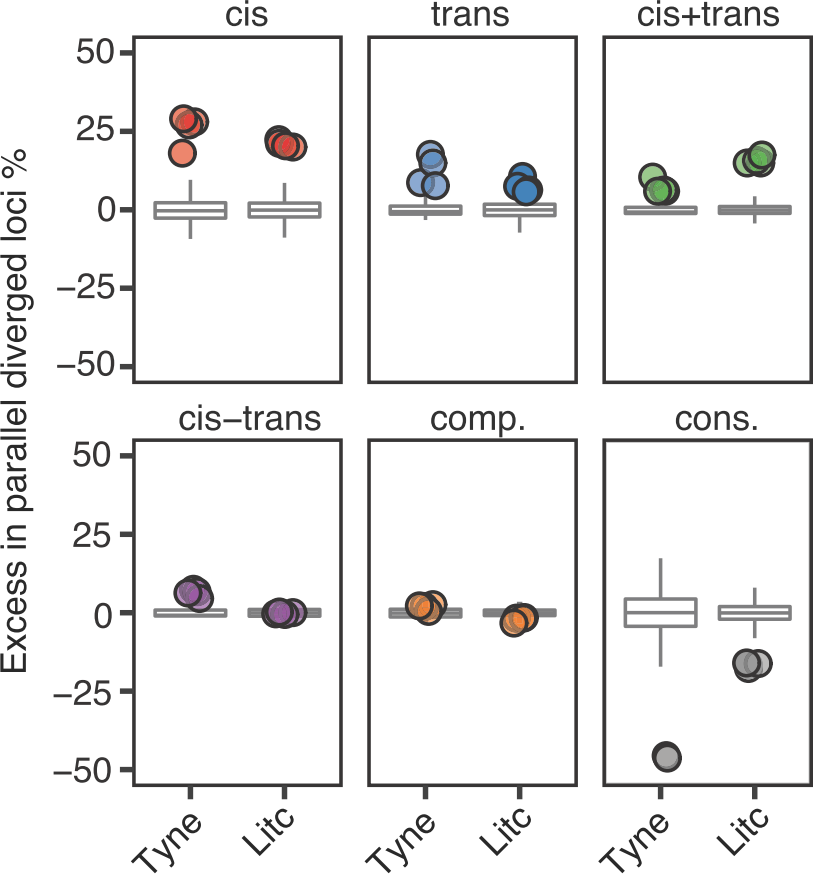
Genes with parallel marine-freshwater divergence in expression are enriched in *cis*-regulation. Overrepresentation of expression regulation types associated with parallel evolving transcripts, compared to the overall frequency of each regulatory divergence type. Points represent the observed values in four F1’s per cross. Box-plots represent the random expectation based on 1000 bootstrap samples of the corresponding number of transcripts from the full set of transcripts suitable for allele-specific expression analysis. Horizontal bar in box-plot corresponds to median, box includes 50% and whiskers 99.3% of random expectation.

In parallel diverging transcripts, ecotype-pairs from both rivers further showed a significant excess of *cis*- and *trans*-regulation of expression divergence acting in the same direction (divergence in *cis*+*trans* co-regulation). Theory and experiments predict that directional selection on expression levels favors the accumulation of *cis*- and *trans*-regulatory changes that have the same direction of effect and act in a cooperative manner towards greater expression divergence (Fraser et al., 2010; Orr, 1998). The observed excess of *cis*+*trans*-regulation in the transcripts that contribute to parallel expression divergence, is consistent with directional selection, as opposed to genetic drift, which should influence amplifying (*cis*+*trans*) and cancelling (*cis*-*trans*) divergence equally (Fraser et al., 2010). Notably, the candidate genes slc12a2, cftr, atp1a1.2, trpv6 and aqp3a that are involved in teleost ion homeostasis were among the parallel diverged transcripts influenced by *cis*+*trans*-acting regulatory divergence.

We next explored the extent of parallelism in the strength (magnitude) of *cis*- and *trans*-regulation among the four populations and stratified loci by their degree of contribution to parallel expression divergence (composite PC loading). For each F1 individual at each locus we quantified *cis-*regulation following Wittkopp et al (Wittkopp, Haerum, & Clark, 2008) as the F1 allele-specific expression ratio, and from there, quantified *trans*-regulation by subtracting the F1 allele-specific expression ratio from the parental expression ratio (Wittkopp et al., 2008). For each river system we then averaged the *cis-*regulatory and *trans-*regulatory values for each locus across each of the four offspring to obtain a measure of the mean degree of *cis-* and *trans-*regulation per locus. We analysed this data with a sliding cut-off, which results in loci with nested extremes of contribution to parallel marine-freshwater expression divergence as measured by the loadings on the composite principal component axis. Overall we see high correlations between river systems in the degree of *cis*- and *trans*-regulation of loci with parallel expression divergence (left and right ends of composite PC axis). However, the pattern is not symmetrical. Parallel divergent loci that are upregulated in freshwater show strong positive correlations among populations in their quantitative extent of both *cis-* and *trans-*regulation (Fig 5). This correlation is lost in loci that do not contribute to parallel expression divergence (composite PC loadings close to zero) and is less consistent among parallel divergent loci downregulated in freshwater fish. We then estimated the mean effect size (magnitude) of *cis*- and *trans*-regulatory effects for subsets of loci and represented them in bins along the parallel expression divergence (composite PC loading) axis. Again, we found that the magnitude of *cis*- and *trans*-regulatory effects is not symmetrical: divergently expressed genes upregulated in freshwater fish show greater effect size in both *cis*- and *trans*-components than their freshwater downregulated counterparts (Fig 5c, Fig S14).

**Figure 5.**
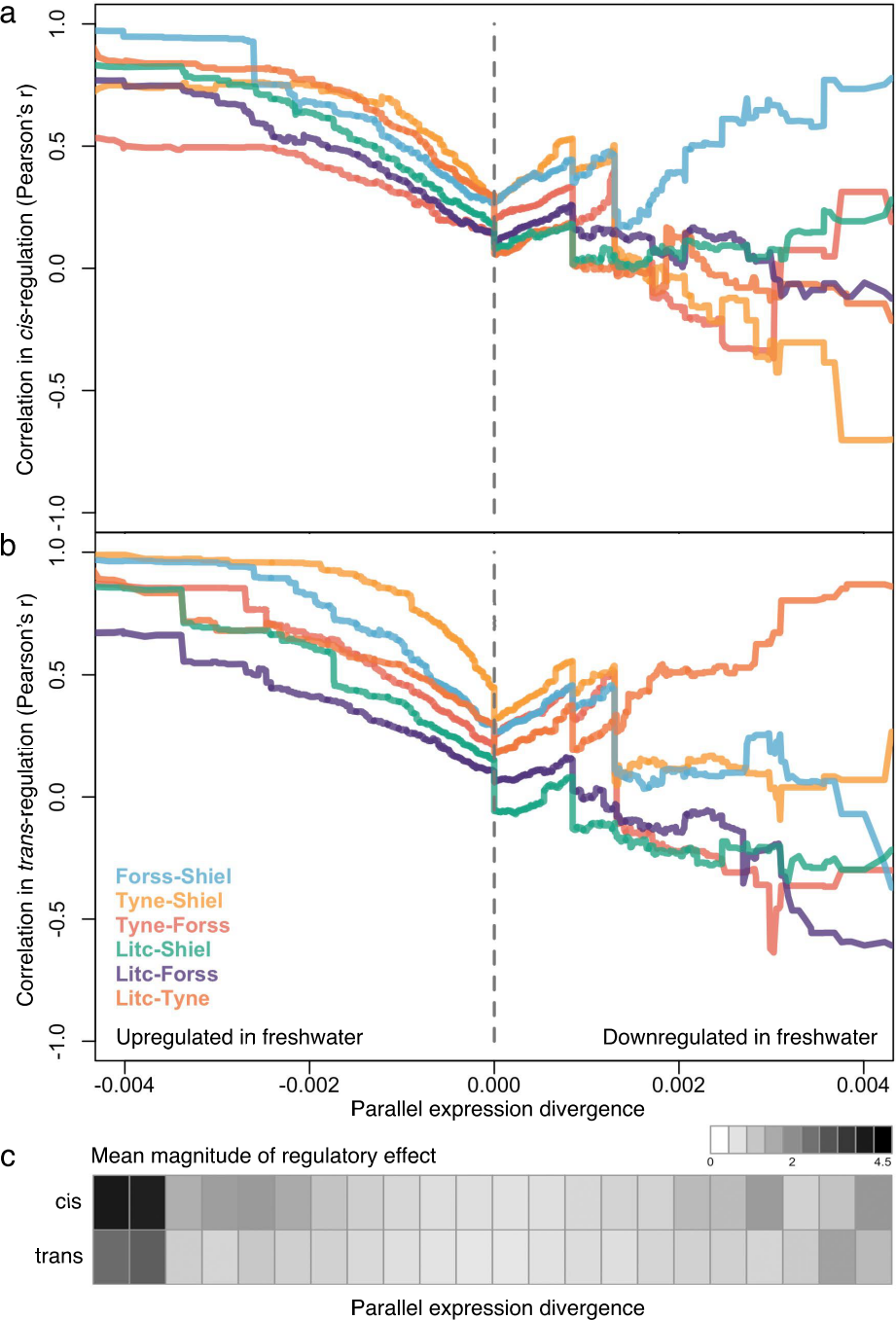
The degree of parallelism (correlation) in the magnitude of cis- (**a**) and trans-regulation (**b**) of marine-freshwater expression divergence among different river systems at loci with increasingly parallel divergent expression (composite PC loadings). Parallel divergent genes that are upregulated in freshwater (strong negative loading on composite PC) show high correlation in their degree of *cis-* and *trans-* regulation among populations. In contrast, loci that have not diverged in parallel (PC loadings close to zero) and loci that are parallel downregulated in freshwater are not highly correlated in the magnitude of *cis-* and *trans-* regulation. For each population pair, Pearson correlation coefficients, r, were calculated for subsets of loci defined by an increasingly extreme positive or negative threshold on the composite PC loading scores. (**c**) The mean absolute magnitude of cis- and trans-regulatory effect across populations in subsets of loci defined by bins of parallel expression divergence (composite PC). For both cis- and trans-regulation higher magnitude effects (darker grey shades) are seen at parallel divergent loci that are upregulated in freshwater. Means and standard errors of effect size per population are shown in Fig S15.

### *Cis*-regulatory effects are additive and insensitive to genetic and environmental context

In order to evaluate the possible factors that may facilitate *cis*-regulation as a predominant source of parallel regulatory differences, we investigated: the mode of inheritance of different regulatory classes; the consistency of regulatory mechanisms in different genetic backgrounds; and the consistency of regulatory mechanisms in different water salinities (a major environmental contrast between freshwater and marine ecosystems).

The mode of inheritance of a given trait, including expression traits, may have a direct impact on the rate of adaptation (Lemos et al., 2008). Among the modes of inheritance, additivity is associated with the greatest efficiency of bi-directional selection, because the contribution of each allele can be seen by selection (Hartl & Clark, 1997). To investigate this, we quantified the dominance/additivity ratio following Gibson et al (Gibson et al., 2004) by taking the ratio between the mid-parental normalized read counts and the deviation from this mid-point in F1 hybrids. *Cis-*regulatory divergence showed the strongest level of additivity (Fig 6a), consistent with evolutionary potential for fast allele-frequency changes at *cis*-regulatory elements. These findings are consistent with previous studies showing that *cis-*divergence is linked to higher additivity of between-species expression differences (Lemos et al., 2008; McManus et al., 2010), except that here, we observed such divergence already among marine-freshwater ecotypes pairs with on-going gene flow, highlighting the role of *cis*-regulation in adaptive divergence even in nascent population pairs.

**Figure 6.**
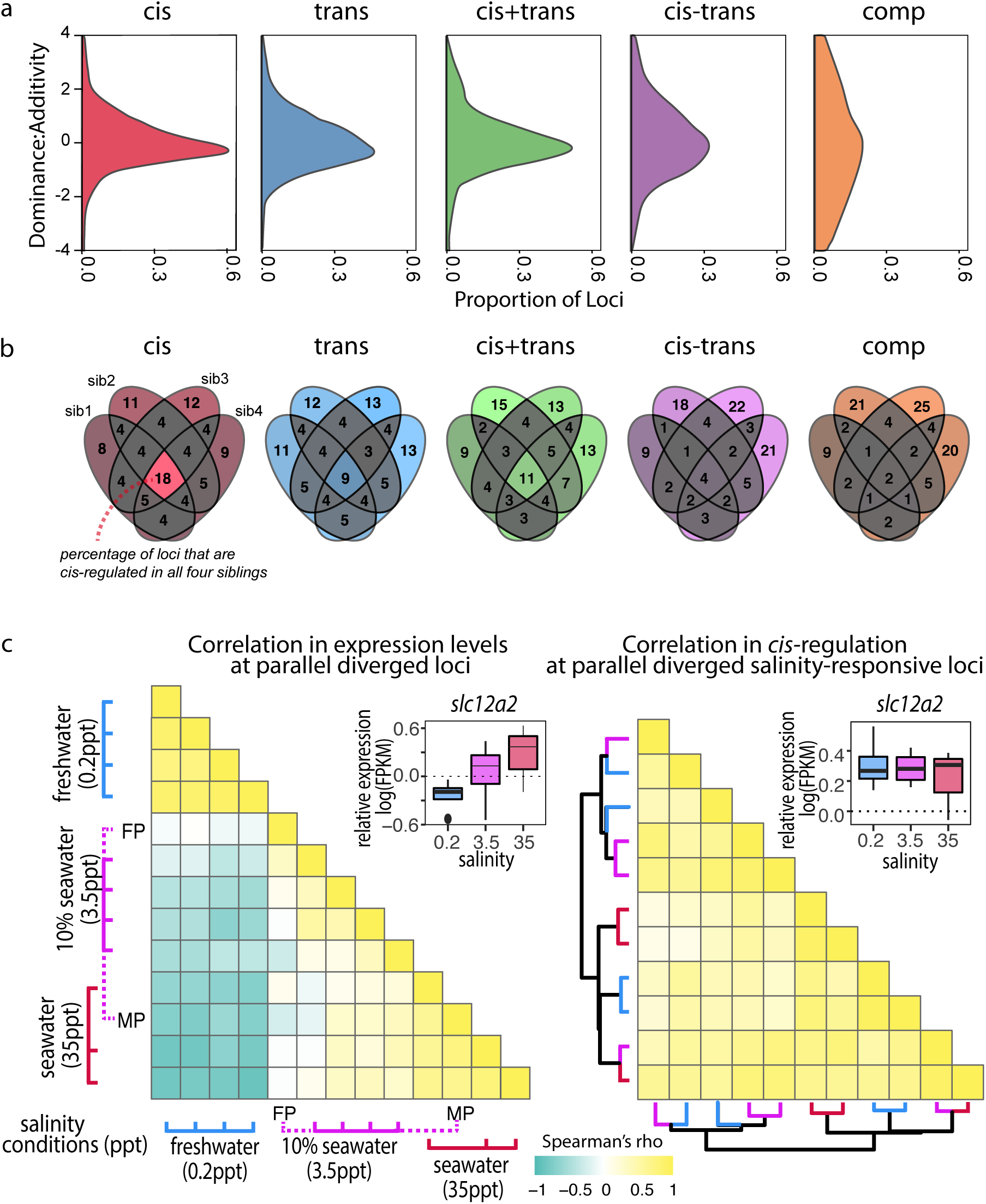
*Cis*-regulatory effects are additive and insensitive to genetic and environmental contexts compared to other regulatory classes, (**a**) Density estimates of dominance-additivity ratio of each locus assigned to the respective regulatory classes (Little Campbell). A ratio of zero indicates additivity, ±1 full dominance, and values >1 or <-1 imply over- or underdominance. The slight bias towards negative values indicates tendency of stronger dominance of low-expressed allele (irrespective of ecotype). (**b**) Sharing of regulatory divergence between siblings as a proxy for the level of epistasis associated with each regulatory class in Little Campbell. The type of regulation observed for each gene is compared among siblings (overlapping ovals) with numbers representing percent of loci shared between siblings (rounded to integer). Color scale ranges from grey (low) to colored (high). *Cis*-regulatory divergence tended to be most stable across genetic backgrounds indicating that *cis*-acting divergence is least influenced by epistasis. (see also data for River Tyne shown in supplement figure S13). (**c**) While the expression levels of parallel diverged transcripts are sensitive to water salinity (left triangle) the degree of *cis*-regulation of these loci is insensitive to water salinity (right triangle). Pairwise Spearman’s correlation (*rho*) in expression levels (left triangle) and allele-specific-expression ratio (right triangle) of parallel diverged loci in marine x freshwater F1 siblings acclimated to different salinity conditions. A negative correlation (cyand) indicates salinity-responsive genes were up-regulated (or show allele-specific-expression bias) in one salinity and down-regulated (or show allele-specific-expression bias in the opposite direction) in another, while a positive correlation (yellow) indicates that expression profiles and allele-specific expression profiles were similar between individuals. Expression levels of an example locus, slc12a2, are shown as an inset above each heatmap. slc12a2 expression increases under seawater conditions, but *cis-*regulatory control of this expression is stable across different salinities. Rows and columns of the right heatmap are ordered based on euclidean distance and the dendogram shows that *cis*-regulation does not cluster by salinity. Colors of dendrogram branches refer to salinity in parts per thousand (0.2-blue, 3.5-pink, 35ppt-red). MP, FP refer to marine and freshwater parents.

Adaptation is expected to favor non-epistatic alleles that confer a stable phenotype irrespective of the genetic background (Stern & Orgogozo, 2009). By using first generation wild derived parents in the crosses for this experiment we have maintained meaningful levels of genetic variation similar to those present in natural populations. The unique combination of parental genetic variation passed on to each F1 offspring enables us to explore the stability of gene expression regulation under different genetic backgrounds. We hypothesised that non-epistatic regulatory divergence should be consistent across siblings due to tight linkage, while epistatic regulatory divergence will leave a pattern of low correlation within the classes of regulatory divergence among F1 siblings. To quantify the level of epistasis in different classes of regulatory divergence we compared the reproducibility of regulatory divergence across siblings of the same F1 family. Our results, consistent in both Tyne and Little Campbell crosses, show that *cis-*regulatory divergence tends to be most stable across genetic backgrounds (Fig 6b and Fig S15) making it a favourable substrate for adaptive divergence.

We next explored the stability in gene regulation under different environmental salinity conditions. Previous studies have reported changes in gene expression plasticity in association with freshwater adaptation (Gibbons et al., 2017; Velotta et al., 2016). We analyzed adult gill transcriptomic and *cis*-regulatory responses to water salinity using full-sib F1 hybrids raised under three different salinity conditions (0.2 ppt (freshwater), 3.5 ppt (approximately equivalent to 10% sea water) and 35 ppt (marine salinity, see methods for more information)

A total of 2542 transcripts were differentially expressed between at least two salinity treatments (FDR 1%), consistent with major changes in gill structure and function upon salinity acclimation (Hwang, Lee, & Lin, 2011). Differences between salinity treatments were mostly driven by freshwater acclimation (0.2 ppt), which elicited a response markedly different from both standard husbandry conditions (3.5 ppt) and typical marine salinity (35 ppt) (Fig 6c, Fig S16). We observed that an unexpectedly high proportion of parallel diverged transcripts also showed an effect of salinity on expression level (97 of 586, hypergeometric test *P*<4.3e-6), suggesting a large proportion of the parallel diverged transcripts in the gill are involved in physiological responses to water salinity.

Plasticity in expression of a given genotype in different environments can be mediated by either *cis* or *trans*-regulatory mechanisms. Focusing on the 97 transcripts with parallel expression divergence that showed a salinity response, we asked whether the regulatory control of these loci is sensitive to salinity conditions. We first calculated an expression profile for each F1 sibling by scaling the FPKM expression of each transcript to the average expression level across all individuals and then compared the profiles using Spearman correlation to capture the strength and direction of correlation in expression profiles among salinity treatments. F1’s raised in similar salinities tended to have high positive correlation coefficients while the expression profiles of individuals raised in freshwater tended to be negatively correlated (opposite expression) with expression in individuals raised in salt (Fig 6c, left triangle).

Then, using allele-specific expression as a proxy for *cis*-regulation we investigated the stability of *cis*-regulation of expression across the salinity treatments. Despite the observed plastic response of gene expression (Fig 6c, left triangle), the degree of *cis*-regulation of salinity-responsive genes with parallel expression divergence was highly correlated across salinities (Fig 6c, right triangle), indicating that the observed salinity response was not caused by *cis*-regulatory elements but largely due to *trans*-acting regulation that influenced the expression of both alleles in similar magnitude. Based on these results we hypothesize that *cis*-regulatory changes provide a mechanism for genetic assimilation of plastic responses into heritable variation (Waddington, 1942) where the effects of regulatory alleles are independent of the environment.

## Discussion

Regulation of gene expression is thought to play a major role in adaptation, yet relatively little is known about the patterns and predictability of adaptive regulatory mechanisms in the early stages of intra-specific adaptive divergence that evolve in the face of on going gene-flow. We characterised parallel expression divergence in the threespine stickleback gill transcriptome - an organ with important respiratory and osmoregulatory functions - and dissected the *cis*- and *trans*-regulatory architecture in four marine-freshwater ecotype pairs in order to infer the rules and patterns shaping genome evolution and influencing rapid adaptation in natural populations.

We found that when sticklebacks are reared under the same standard husbandry conditions, parallel transcriptomic divergence involves only a few hundred genes each with relatively small effect on expression divergence, similar to the small proportion of the genome that shows parallel adaptive divergence at the DNA sequence level (Jones et al., 2012). Parallel divergently expressed genes are proximal to adaptive loci identified from signatures of parallel divergence at the sequence level, and support a role for the reuse of ancient standing genetic variation in the parallel adaptive divergence of gene expression. We also observed a large proportion of loci with marine-freshwater divergent expression that was unique to a local river system (see Supplementary note) indicating there is room for drift and/or local adaptation to play a major role shaping the evolution of divergent gene regulation and gene expression in any given river system. The transcripts showing strongest parallel expression divergence are associated with genes involved in gill ion transport and osmoregulation, a key physiological trait in marine-freshwater divergence, and in agreement with studies in other fish (Velotta et al., 2016). Through population genomic analysis we show how natural selection on divergently expressed genes is shaping the evolution of the genome - strong molecular signatures of selection (elevated FST, reduced Pi) were detected around the transcription start sites of parallel divergently expressed genes.

We found overwhelming evidence for the importance of *cis*-regulation of marine-freshwater divergent gene expression through analysis of allele-specific expression in F1 hybrids relative to their marine and freshwater parents. The predominance of *cis*-regulation was observed in marine-freshwater ecotypes from four independent river systems, in analysis of all informative transcripts genome wide, and enriched in the set of loci identified as having evolved parallel marine-freshwater divergence in expression across rivers

In animals and yeast, *cis*-regulatory differences contribute strongly to divergence in gene regulation over long evolutionary divergence scales (Coolon et al., 2014; Emerson et al., 2010; Lemos et al., 2008; Prud’homme, Gompel, & Carroll, 2007; Stern & Orgogozo, 2009). The stickleback ecotype pairs studied here diverged from each other within the last 10-20k years (approximately 10-20k generations) following the retreat of the Pleistocene ice sheet, indicating that strong *cis*-regulatory divergence can also evolve in relatively short timescales. While adaptation via re-use of ancient standing genetic variation is important in sticklebacks and may explain some part of the predominance of *cis*-regulatory changes, we also found that the extent of genomic divergence varied substantially between the parallel evolving stickleback ecotype-pairs we studied, and the proportion of *cis*-regulatory divergence scaled positively with this genomic divergence. This suggests that much of the genetic differences that accumulate in the early stages of adaptive divergence with gene flow translate to *cis-*regulatory differences. Mutation-accumulation experiments have shown that genetic drift, which promotes random regulatory changes, is biased towards *trans-*acting divergence due to a larger *trans*-mutational target size (Landry, Lemos, Rifkin, Dickinson, & Hartl, 2007b). Since most expression changes between freshwater and marine ecotypes however tended to be regulated in *cis-*, this points towards a stronger contribution of selection rather than drift.

Our results further indicate that reinforcing *cis* and *trans-*acting regulatory variation that act in a manner to amplify one another has an important contribution to early ecotype divergence with gene flow. Overrepresentation in *cis+trans*-coevolution seemed to grow with increasing genetic and expression divergence between lineages, which is notable in the context of incipient ecological speciation. Given enough evolutionary time, the accumulation of amplifying *cis+trans-*regulatory divergence independently in diverging populations may lead to the evolution of genetic (Bateson-Dobzhansky-Muller) incompatibilities. Because recombination between coevolved *cis-* and *trans-*regulatory factors disrupts their combined phenotypic effects, it is expected that selection would favor linkage disequilibrium between the coevolved regulatory factors (Verta, Landry, & Mackay, 2016). Association between increasing *cis+trans*-regulation and genome-wide divergence as seen here suggests that selection against recombination between coevolved regulatory factors may contribute to increased genetic divergence as adaptation proceeds and thus shape the genomic landscape of incipient speciation.

Parallel adaptive divergence of marine and freshwater ecotypes provides biological replicates of the evolutionary process which we can interogate to identify common patterns governing the molecular basis of adaptation. We not only identified parallelism across marine-freshwater ecotype pairs in the predominance of *cis*-regulation of divergent gene expression, but also parallelism in the quantitative extent of both *cis*- and *trans*-regulation of divergently expressed loci. This strong parallelism was particularly notable in parallel divergently expressed genes that are upregulated in freshwater, and considerably less strong and less predictable in parallel divergently expressed genes that are downregulated in freshwater. Similar to the underlying shared genetic basis of freshwater adaptation due to the parallel reuse of standing genetic variation, our results suggest we can not only predict that freshwater populations are likely to carry the same alleles at adaptive loci across the genome, but also that the magnitude and extent of *cis*- and *trans*-upregulation of divergently expressed genes is likely to be shared among freshwater populations. It is possible that there are fewer molecular mechanisms available to freshwater fish to ‘turn on’ gene expression and more diverse mechanisms to ‘turn-off’ gene expression. Coinciding with the stronger parallelism in genes upregulated in freshwater we observed a tendency for the *cis*- and *trans*-regulatory effect size to be of larger magnitude than the *cis*- and *trans*-regulatory effects of genes downregulated in freshwater. It is plausible that this larger effect size enables selection to be more efficient and contributes to the observed stronger parallelism. Further, the marine ecotype is thought to be the ancestral state in threespine sticklebacks and has been evolving in a comparatively stable marine environment for millions of years. Under this evolutionary context, freshwater populations have a comparatively smaller effective population size with reduced access to pre-existing adaptive standing genetic variation, and have been subject to more recent and potentially stronger selection pressures than their marine ecotype counterparts. Thus marine ecotypes have more opportunity to evolve via soft sweeps on standing genetic variation. Combined this evolutionary context is likely to have influenced which molecular mechanisms of gene expression regulation are visible, and efficiently respond to selection, resulting in relatively few paths to evolve adaptive upregulation of gene expression in freshwater populations compared to marine.

We note that our results contrast with a recent study by Hart et al (Hart et al., 2018) who reported a predominance of, and parallelism in the *trans*-regulatory control of marine-freshwater divergent gene expression in the pharyngeal tooth plate. One possible biological explanation for this might be differences in the multifunctional (pleiotropic) functional roles of the gill and its likely complex genetic architecture compared to dental tissue with a less pleiotropic functional role and more simple genetic architecture involving a few large effect loci upstream of other factors (Cleves et al., 2014) that cause a dominant *trans*-regulatory signal in the allele-specific expression assay.

Our results indicate that parallel evolving divergence may converge on *cis-*regulation driven in part by a higher level of additivity and lower level of epistasis of *cis-*regulatory factors. Known as the effect of “*Haldane’s sieve*”, the fixation of beneficial recessive alleles are hindered due to its phenotype being hidden from selection (Haldane, 1927). When alternative alleles are favored in their respective environments, as seem to be the case in sticklebacks, additive alleles have the highest likelihood of becoming fixed in both populations as both alleles can rise quickly to a high frequency. *Cis-*regulatory variation also tended to have a lower level of epistasis within populations compared to other types of regulatory effects. Epistatic regulatory alleles tend to have different phenotypic effects depending on the genetic background, therefore inducing unpredictable fluctuation in expression levels. Hence, parallel evolution of gene expression seems to favor non-epistatic regulatory alleles that have similar effects on expression levels independently of the genetic background.

In addition to promoting divergence in expression levels that is independent of genetic backgrounds, adaptation-with-gene-flow is predicted to promote reduced expression plasticity, especially in cases where selection is strong (Stern & Orgogozo, 2009). Towards this end, our results indicate that *cis*-regulation may play a particularly important role in the evolution of reduced plasticity. Since plasticity is known to be involved in the divergent adaptation of marine and freshwater sticklebacks we investigated the sensitivity of gene expression and its *cis*- and *trans-*regulation to environmental salinity. Genetic assimilation involves the heritable encoding and loss of plasticity of a once plastic trait (Waddington, 1942). Previous studies have found evidence for genetic assimilation in gill gene expression evolution between stickleback ecotypes (McCairns & Bernatchez, 2010). While we found a strong component of plasticity in gene expression among siblings raised in different salinities, the *cis*-regulation of this gene expression was stable and insensitive to differences in the environmental conditions. From this we infer that the observed plasticity in expression is likely mediated via *trans*-regulation, and hypothesize that the stable *cis*-regulatory component may serve as a mechanism for genetic assimilation.

The importance of *cis*-regulation underlying parallel marine-freshwater expression divergence has implications for our understanding of genome function in natural populations in the early stages of adaptive divergence. The predominantly *cis-*regulated, parallel diverged loci are dispersed throughout the genome but proximal to previously identified adaptive loci. We observed signatures of selection around their transcription start sites likely to be a result of natural selection acting on *cis-*regulatory elements. We show that *cis-*regulation acts in an additive manner, and is robust to both different environmental contexts, and potential epistasis caused by differences in genetic background. Comparing across marine-freshwater ecotype pairs from independent river systems, we observed parallelism and large magnitude effect sizes for *cis*-regulation of divergently expressed loci up-regulated in freshwater. These features make *cis-*regulation a well-poised target for natural selection and may explain parallelism in the predominance and quantitative extent of *cis*-regulation in the early stages intraspecific adaptive divergence. Combined our study highlights how natural selection on adaptive *cis-*regulation is a likely contributor to the dispersed genomic landscape of adaptation in sticklebacks.

## Data access

Data will be deposited to the Sequence Read Archive following manuscript acceptance. All scripts used in data analysis will be made available at https://github.com/jpverta/ following manuscript acceptance.

## Supporting information

## Acknowledgements

The authors acknowledge the Semiahmoo First Nation and Dolph Schluter for collaboration with sampling Little Campbell River sticklebacks and Saad Arif, Cecilia Martinez, Frank Chan, Paloma Medina and Li Ying Tan with stickleback sampling. We thank Stanley Neufeld, Enni Harjunmaa, Muhua Wang and Frank Chan for helpful comments on the manuscript and Vrinda Venu for assistance in DNA library preparation and sequencing. FCJ is funded by ERC Consolidator Grant, Deutsche Forschungsgemeinschaft and the Max Planck Society.

## Author contribution

JPV and FCJ designed the study. JPV conducted the experiments and analyzed the data. FCJ contributed to data analysis. JPV and FCJ wrote the manuscript.

## Materials and Methods

### Sample material

We captured freshwater-resident and anadromous marine sticklebacks from Little Campbell River, Canada, and Rivers Tyne, Forss and Shiel, Scotland, with wire-mesh minnow traps. Sampling locations are given in table S1. We identified freshwater resident and anadromous marine ecotypes based on their lateral plates. Most freshwater-resident populations of sticklebacks are plate-less, whereas anadromous marine forms exhibit complete lateral plating (Bell & Foster, 1994). Consistently, in most cases the large majority of fish captured in freshwater were plate-less and anadromous marine fish captured near the mouth of the river/lake were completely plated. We generated within-ecotype crosses via *in vitro* fertilisation of gravid females with males within ecotypes and transported the fertilized eggs to a common-garden environment at the Max-Planck campus in Tübingen in reverse-osmosis water supplemented with Instant Ocean salt to 3.5 ppt (~10% sea water salinity). The Max Planck Society holds neccessary permits to capture and raise sticklebacks. All animal experiments were done in accordance to EU and state legislation and avoiding unneccessary harm to animals.

The fish strains were raised in individual 100 liter tanks with 40-50 fish per tank in 3.5 ppt salinity and alternating light cycle of 16 hour light and 8 hour darkness of 6 months. No significant mortality occurred during transfer or captivity. Fish were raised with a diet of fry (freshly hatched artemia), juvenile (artemia Daphnia and cyclops) and adult food (bloodworm, white mosquito larvae, artemia, mysis shrimp and Daphnia) until adults. We selected four unrelated fish from independent field crosses per ecotype from Little Campbell River and River Tyne strains, and one fish per ecotype from Shiel and Forss strains. Exception to this was one Tyne freshwater male fish that was the progeny of unrelated lab-raised freshwater parents, which was included to complete the sampling. We in-vitro crossed one marine female and one freshwater male for each strain, and raised the F1 individuals in identical conditions (each cross in individual tank) as the parents until they were reproductively mature.

For the Tyne cross, we separated the F1 clutch into three at 3 months of age and transferred the F1s into separate 100 liter tanks with 3.5 ppt water. We added either 0.2 ppt or 35 ppt water in increments of 20 liters at a time twice a week over the course of one month to acclimate fish in two of the tanks to different water salinities. After one month of acclimation all water in the two tanks was changed to either 0.2 or 35 ppt. We raised the fish in 0.2, 3.5 or 35 ppt water for additional three months before harvesting tissue.

### Sample preparation and RNA-sequencing

We harvested gill tissue for all strains and F1’s, all staged as adults and reproductively active (gravid females and males exhibiting mating coloring). We flash-froze gills on liquid nitrogen and stored in −80 degrees C until used for mRNA-extraction. We disrupted gill tissue with a pestle on liquid nitrogen and extracted mRNA using Dynabeads mRNA direct kit (Invitrogen) and following manufacturer’s instructions, followed by DNase treatment with the Turbo DNA-free kit (Ambion). We verified mRNA quality using Agilent BioAnalyzer.

We used 150 nanogram of mRNA to construct strand-specific RNA-seq libraries using the TruSeq Stranded RNA-seq kit (Illumina). We verified library yield using Qubit and size distribution using BioAnalyzer. We optimized library construction protocol to result in mRNA insert size distribution centered on 290 base pairs. We pooled the libraries in equimolar amounts and sequenced in pools of eight samples on a HiSeq-3000 instrument, producing 150 bp paired-end reads (Table S2). We included replicate sequencing libraries in different lanes of the same run and different runs of the same instrument in order to measure the effect of batch on final data (none observed).

### Gill transcriptome assembly from RNA sequencing

We verified read quality with *FastQC* software (http://www.bioinformatics.babraham.ac.uk/projects/fastqc/) and trimmed the reads of sequencing adapters using TrimGalore (http://www.bioinformatics.babraham.ac.uk/projects/trim_galore/).

We aligned RNA-seq reads from pure strains to the UCSC stickleback genome reference (“gasAcu1”) with *STAR* aligner (Dobin et al., 2013). We opted for running *STAR* in two-pass mode, gathering novel splice junctions from all pure-strain samples for the second alignment pass. After experimenting with alternative parameters, we opted for the following: –*outFilterIntronMotifs* RemoveNoncanonicalUnannotated – *chimSegmentMin* 50 –*alignSJDBoverhangMin* 1 –*alignIntronMin* 20 –*alignIntronMax* 200000 –*alignMatesGapMax* 200000 –*limitSjdbInsertNsj* 2000000.

We followed the *Cufflinks2.2* pipeline (Trapnell et al., 2012) for reference-guided transcriptome assembly and transcript and isoform expression level testing. We used *Cufflinks2* to assemble aligned RNA-seq reads into transcripts, using Ensembl gene models for stickleback (version 90) as guide and the following parameters: –*frag-bias-correct* gasAcu1.fa –*multi-read-correct* –*min-isoform-fraction* 0.15 –*min-frags-per-transfrag* 20 –*max-multiread-fraction* 0.5. We produced a single merged transcriptome assembly based on all pure strains using *CuffMerge* and used this in all subsequent analyses for all samples.

Finally, we used *CuffDiff* to test expression differences between male and female fish from freshwater and marine ecotypes of Tyne and Little Campbell river (combined). Transcripts with sex-dependent expression at FDR 10% (N=278) were excluded from analysis of genetic architecture of expression divergence (see below), but included in all other analyses where the number of male and female fish were balanced across the experimental contrast.

### Principal Component Analysis

We summarized read counts over transcript models using the *Cufflinks* function *CuffQuant with ‘fr-firststrand’ strand-specific RNAseq library type and other settings as default* and normalized read counts to total library sizes using *CuffNorm*. We subsequently transformed the read data with the *DESeq2* (Love, Huber, & Anders, 2014) function *varianceStabilizingTransformation* so that the variance in read counts was independent of the mean, following the steps outlined in the *DESeq2* manual. We used the *R* (Ihaka & Gentleman, 1996) function *prcomp* with the option *scale=FALSE* to calculate PCA on expression level co-variances using data from all transcripts. We calculated the PCA based on a balanced set of four freshwater and four marine ecotypes from both Tyne and Little Campbell, and projected the single ecotypes from Forss and Shiel to principal components 2 and 5 using the *R* function *scale*. We verified the absence of batch effects in the RNA-seq data by PCA analysis of a replicate sequencing library sequenced on different lanes of the same run and on different runs. Technical variation was much smaller than biological variation.

As described in the main text, a combination of principal components 2 and 5 best described freshwater-marine divergence in transcriptomes in our dataset. We therefore defined a composite principal component by summing principal components 2 and 5, weighing each with the percentage of variation explained. Finally, we extracted principal component 2 and 5 loadings for each transcript and used the identical approach to calculate transcript loadings on the composite principal component. This procedure produced a loading value for each transcript that described the importance of that transcript in parallel freshwater-marine expression divergence.

### Gene Ontology analysis

Tests for enrichment of genes involved in specific biological processes, molecular functions and cellular components among top ranking differentially expressed genes was performed using GOrilla (Eden, Navon, Steinfeld, Lipson, & Yakhini, 2009). Genes were sorted by CuffDiff differential expression q-values or by composite PC loading score and, because stickleback genes are largely unannotated for gene ontologies, were mapped to mouse orthologs (the vertebrate with the highest GO annotation quality (ref: https://www.ncbi.nlm.nih.gov/pmc/articles/PMC2241866/) using a REST command to access the Ensembl precomputed ortholog database. We tested for significant enrichment of gene ontologies (biological processes, molecular functions, and cellular compartments) with p-values less than 1×10^−5^. A similar approach using both human and zebrafish orthologs revealed enrichment in many of the same gene ontologies (not shown).

### DNA preparation and sequencing

We extracted genomic DNA from fin clips of six unrelated fish per ecotype (six females of each ecotype from Little Campbell, three males and three females of each ecotype from Tyne) using standard lysis buffer and proteinase-K digestion, followed by SPRI bead extraction with Ambion magnetic beads. We verified DNA quality on agarose gel and quantified DNA concentration using Qubit.

We fragmented 700 ng of gDNA with Covaris instrument and selected 300-500 bp DNA fragments for library construction using double-sided SPRI selection. We constructed DNA-sequencing libraries using a custom protocol that includes DNA end-repair, A-tailing and Illumina TruSeq adapter ligation, followed by 6 cycles of PCR amplification. We verified library fragment size using BioAnalyzer and quantified library concentrations using Qubit. Libraries were sequenced with Illumina HiSeq-3000 instrument to an estimated whole-genome coverage of 10-40X (Table S5).

### DNA-sequencing read processing, alignment and SNP discovery

We verified DNA-sequencing read quality using FastQC and trimmed adapter sequences using TrimGalore. We aligned DNA-seq reads to the stickleback reference genome sequence (“BROAD S1” Jones et al., 2012) using *BWA mem* (Li, 2013).

We used the Broad Institute Genome Analysis Tool Kit (*GATK*) (McKenna et al., 2010) to call Single Nucleotide Polymorphisms (SNPs) in genomic resequencing data, following the DNA-seq best practices (as in June 2016). We ran *GATK HaplotypeCaller* individually for each sample and defining parameters *-stand_call_conf* 30 *-stand_emit_conf* 10 *–emitRefConfidence* GVCF *-variant_index_type* LINEAR *-variant_index_parameter* 128000. The step was followed by joint genotyping using *GenotypeGVCFs* after which we excluded indels from the analysis. The final step was *VariantQualityScoreRecalibration* (*VQSR*). Samples from each ecotype-pair as well as each controlled cross were analyzed together (but separately from other ecotype-pairs or controlled crosses) from *GenotypeGVCFs -* step onwards. Because stickleback lacks a set of known variant sites, we opted for using a hard-filtered set of SNPs as “true” set of SNPs (with *prior=*10). We used the *GATK SelectVariants* tool to extract a training set that fulfilled the following thresholds: *QD*>30, *FS*<60, *MQ*>40, *MQRankSum*>-12.5 and *ReadPosRankSum*>-8. After inspection of *VQSR* tranche plots, we selected the 99.9% quality tranche for downstream analysis, which captured 1.66M and 3.52M SNPs with transition/transversion ratios of 1.14 and 1.18 for Tyne and Little Campbell population genomic analyses respectively, and 2.27M, 1.62M, 2.75M, 1.54M SNPs with transitition/transversion ratios ranging from 1.16-1.17 for parents of the four crosses (strains from Forss, Tyne, Little Campbell, Shiel) used in allele-specific expression analysis.

### Population genetic analyses

Population genetic analyses were based on a set of six freshwater and six marine fish from both Tyne and Little Campbell River. From the GATK variant calling analysis (described above) we identified over 3.5 million SNPs in the Little Campbell populations and over 1.6 million SNPs in Tyne. We calculated per-site statistics for Weir & Cockerham’s FST (Weir & Cockerham, 1984) and average pairwise-nucleotide diversity (Pi) genome-wide, and for 400 kilobase (kb) regions centered on transcription start sites (TSSs) using *VCFtools* (version 0.1.14) (Danecek et al., 2011) allowing for a maximum of 4 missing genotypes per SNP for calculation of FST and a maximum of 2 missing genotypes per SNP for calculation of Pi, corresponding to a maximum of 20 % missing genotypes in each case. Negative F^ST^ values were rounded to zero. CSS score was calculated based on Pi and following (Jones et al., 2012) in 10kb non-overlapping windows across the genome. We used the 10kb windows to assign genome regions as having strongest level of parallel genetic divergence between Tyne and Little Campbell freshwater and marine ecotypes, keeping the top 1% windows with the highest CSS score.

We used custom R scripts to calculate F_ST_ and Pi in 1kb windows centered on transcription start sites (TSSs, as reported by *CuffLinks*). We used custom R scripts and the R package *GenomicRanges* to compare the genomic cordinates of transcripts showing parallel expression divergence to the cordinates of the genomic windows showing high CSS values. We calculated the average distance between CSS outlier windows and transcripts in increments of 10kb and compared the average distance to distances calculated based on 1000 randomized sets of transcripts.

### Allele-Specific Expression analysis

We defined a set of high-confidence SNPs for Allele-Specific Expression (ASE) analysis, based on genomic resequencing of parent fish used in controlled crosses (above). We then used *GATK FastaAlternateReferenceMaker* to mask the stickleback reference genome in the corresponding position with N’s in order to avoid preferential mapping of reference SNP alleles. We aligned RNA-seq reads from each F1 and the parents onto the N-masked reference genomes using *STAR* and parameters as described above, with the exception that we allowed for only uniquely mapping reads (–*outFilterMultimapNmax* 1). No significant preferential mapping of reference SNPs was observed after these steps.

We selected SNPs where the genotypes of the parents were covered by at least 10 DNA sequencing reads in each parent and where the genotypes of the parents were homozygous for different alleles. These SNP positions were assigned as informative for allele-specific analysis in each cross. Expression levels for allele-specific analyses were represented as read counts overlapping informative SNP positions. We generated allele-specific read counts for F1’s and parents with the *GATK ASEReadCounter* tool and enabling default filters. We verified that parents had more than 99% counts assigned to right genotypes and excluded the few SNPs where the RNA-seq reads indicated that both alleles were expressed in a parent that should be homozygous. We combined all individuals in each cross (parents and F1’s) in one data frame and normalized read counts between individuals to the total library size using *DESeq2* function *estimateSizeFactors* in order to have equal power across F1s. We then filtered for SNP positions covered by more than 10 reads in at least one F1 in order to avoid underpowered tests of allele-specific expression at loci showing no or very low expression. Finally, we intersected the SNP-based results with transcript models from our reference-guided assembly using the R package *GenomicRanges*, assigning each informative SNP position to an expressed locus and exon.

We tested for ASE in each informative SNP position using a binomial exact test in *R* and an FDR level of 10% (Storey, 2002). Normally the null hypothesis of binomial test is 0.5, in our case meaning that 50% of the RNA-seq reads represented either marine or freshwater parent alleles. Our approach takes into account a possible residual effect of preferential mapping of reference allele reads by calculating the null-hypothesis for the binomial test based on the ratio of all reference reads over all alternative reads per each F1 following (Buil et al., 2015). The null hypothesis calculated this way was between 0.5 and 0.52, indicating that the residual effect of preferential mapping of reference alleles, if detected, was small. Finally the results for the ASE test were converted from reference allele versus alternative allele format into marine parent versus freshwater parent by comparing to the genotypes of the parents.

Following ASE testing we tested for analogous expression difference between parents in the corresponding SNP positions, again using binomial exact test and FDR of 10%. We tested for difference in allele-expression ratio versus parental expression ratio with Fisher’s exact test. We then compared ASE significance, ASE sign and ASE magnitude to parental expression difference in order to dissect parental expression differences into divergence classes following (Landry, Wittkopp, Taubes, Ranz, Clark, & Hartl, 2005), outlined in table S4. We verified that our results were robust towards sequencing depth by down-sampling to 30 million RNA-seq reads per sample (Fig S12).

Given that the median number of assembled transcript isoforms per gene (locus) is 3, and the mean number of ASE informative SNPs tagging a given gene (locus) range from 2.0–6.2, we concluded that the level of evolutionary divergence between marine and freshwater stickleback strains used in our study was insufficient to dissect the cis-vs trans-genetic architecture underlying expression divergence at the transcript (isoform) level (see Supplemental Note for more information). We instead analyzed allele-specific expression at the gene (locus) level, classifying cis-/trans-architecture of each gene into one divergence class based on the SNP that showed the highest statistical support (see above).

We performed concordance analysis for validating the reproducibility of allele expression levels and divergence classes on SNPs assigned to the same exon and the same transcript (Supplemental Note). For the final classification of transcripts to divergence types, we ordered informative SNPs per transcribed locus per F1 by the product of the *P*-values of the three tests (above) and selected the SNP that had the lowest product of *P-*value as a representative SNP for that transcript, per F1. We selected this approach among alternatives after taking into account the high concordance of allele expression levels, relatively complex isoform expression patterns, analysis sensitivity, statistically balanced approach, and parsimony in biological explanation of expression divergence (Supplementary Note).

A small number of SNPs showed monoallelic expression in F1’s where RNA-seq reads overlapping one of the parent alleles were not observed (e.g. 1129 SNPs in Little Campbell River cross). Analysis of RNA-seq read coverage indicated that these SNP positions had lower coverage specifically of the alternative allele, and this effect was not observed when evaluated based on all SNPs or subsets of SNPs e.g. assigned as *cis-*diverged. This indicated that the cases of monoallelic expression are likely caused by mappability issues, and that the issue was specific to SNPs showing monoallelic expression. Although including these SNPs did not impact the results in a significant way we decided to exclude SNPs showing monoallelic expression in F1’s or parents from the analysis.

We summarized the frequencies of transcripts assigned to divergence classes for each F1 using custom R scripts, excluding transcripts that showed differential expression between sexes at FDR 10%. We then assigned transcripts to two classes according to whether or not they showed parallel expression divergence in PCA and differential expression analysis. We tested for over-representation in divergence classes using a randomization test. We draw 1000 sets of 586 randomly chosen transcripts from the whole dataset, calculated the frequencies of divergence classes and compared the random expectation to our set of parallel diverged transcripts.

### Salinity response

A clutch of Tyne marine x freshwater F1 siblings were raised in standard laboratory ppt salinity until 3 months old, separated into three groups and acclimated to over four months. At reproductive maturity we analyzed the gill transcriptomes of four F1’s from each salinity using RNA-seq.

For testing the effect of salinity acclimation on gene expression in F1 gills, we estimated transcript-level expression using *CuffQuant* and normalized counts for each sample to total library sizes using *CuffNorm*. We then imported the gene count tables into *R* and tested for expression differences between salinity treatments using contrasts and an FDR level of 1%, as implemented in *DESeq2*.

For analysis of expression profiles, we imported FPKM values from *CuffLinks* into R. The FPKM values were highly correlated with normalized expression values from *DESeq2 VarianceStabilizingTransformation*, and allow for a more intuitive interpretation. We sub-set the data to only include the transcripts showing differential expression between at least one contrast and parallel expression divergence (N=97). We then log-transformed and normalized the expression of each transcript to the average expression level across all samples to produce an expression profile that represent expression in a given sample relative to others (for that transcript). We then compared the profiles of samples with Spearman correlation. A correlation approaching 1 indicates that expression profiles tended to be similar relative to other samples. In contrast, a correlation approaching −1 indicated that sample profiles were mirror-images of one another. Finally, we clustered the samples based on euclidean distance and visualized the sample similarity profiles using the *pheatmap R* function.

For analysis of allele-specific expression in salinity treatments, we imported allele-specific counts over informative SNPs (as defined above) into R and transformed the counts into log-fold change of marine over freshwater allele. We identified one of the samples acclimated to 35 ppt as having outlier allelic expression levels very different from all other samples and excluded the sample from further analysis. We intersected the SNPs with transcripts that exhibited salinity response and parallel expression divergence (N=11 transcripts). We measured the similarity of fold-change expression differences between alleles across samples with Spearman correlation, analogous to FPKM counts.

## Supplementary Note

### Hierarchical clustering of expression

The gill transcriptome can be characterized by 5 major groups of loci (Fig S2) comprising 766 highly expressed genes (2.6%) showing strong enrichment for biological processes including mitochondrial respiration, ATP synthesis coupled proton transport and cytoplasmic translation, 4337 moderately expressed genes (14.8%) enriched for metabolic processes such as mRNA processing and RNA splicing, and functions such as cadherin mediated cell adhesion, 6950 lowly expressed genes (23.7%) enriched with genes involved in protein modification and chromatin modification processes, 5565 low or partially expressed genes (19.0%) enriched in genes involved in developmental processes and regulation of developmental processes, and 11677 predominantly unexpressed genes (39.9%).

### Defining parallel expression divergence using a parametric test

For parametric testing of parallel expression differences between ecotypes, we combined samples from Little Campbell and Tyne and tested for differential expression using *CuffDiff,* specifying the parameter -*dispersion-method* per-condition. We imported the results tables into *R* and selected transcripts that were tested and where the q-value was less than 0.2. We additionally required that the mean expression difference between freshwater and marine ecotypes was of the same sign in Tyne and Little Campbell.

With a false discovery rate of 20%, we identified 120 loci that showed both significant differences in mean and consistent direction (sign) of divergence between ecotype strains from different river systems. The differentially expressed genes tended to have large composite PC loadings (Fig S5). Top ranking genes include Na-Cl cotransporter (slc12a10), Basolateral Na-K-Cl Symporter (slc12a2), cation proton antiporter 3 (slc9a3.2), Potassium Inwardly-Rectifying Voltage-Gated Channel (kcnj1a.3), potassium voltage-gated channel (KCNA2), Epithelial Calcium Channel 2 (trpv6), Sodium/Potassium-Transporting ATPase (atp1a1.4), aquaporin 3a (aqp3a) — genes known to play a role in osmoregulation in fish and other organisms. This set of differentially expressed loci also include a microRNA (mir-182), 31 loci that are annotated in previous Ensembl gene builds but have unknown function inferred from protein homology to other organisms, and 30 entirely novel loci that have no overlap with gene annotations from Ensembl gene build 90.

We used *CuffDiff* also for testing of ecotype-specific expression divergence specifying the same parameters as above. For this analysis, freshwater-marine expression differences were tested separately for Tyne and Little Campbell and the results were compared using *R*. Transcripts that were differentially expressed in both ecotype-pairs with FDR 20% and where the ecotype difference was of the same sign were assigned as “parallel”, whereas if the signs were opposite the transcripts were assigned as “anti-parallel”.

Overall, 719 differentially expressed transcripts (FDR 20%) were identified using a parametric analysis, the majority of which (N=515) had marine-freshwater differential expression unique to Little Campbell compared to N=157 uniquely differential in Tyne strains. Consistent with largely river system-specific ecotype expression divergence only four percent of loci (N=29) show parallel expression divergence in both rivers (significant differential expression and identical sign of expression difference in both Tyne and Little Campbell) while 2.5% (N=18) show anti-parallel expression divergence (significant differential expression and opposite sign of expression difference).

### CSS based on Tyne and Little Campbell population data

We calculated a Cluster Separation Score (CSS) in 10kb non-overlapping windows across the genome following (Jones et al., 2012). The CSS score reflects parallel genetic divergence between freshwater and marine fish irrespective of their geographic origin. We assigned the genomic windows with the extreme 0.5% CSS values as regions showing the strongest signal of parallel genetic divergence (449 windows). More than 26 percent of the transcripts evolving in parallel between Tyne and Little Campbell were situated within 10kb of regions of parallel genetic divergence (randomization test, *P*<<0.025, Fig S6), which is two times more than what was observed for the global set of regions showing parallel genetic divergence (main text).

### SNP concordance analysis

We performed SNP concordance analysis to validate the reproducibility of allele expression levels between SNPs assigned to same exons and to different exons of the same transcript. Our assumption for this analysis was that individual SNPs assigned to the same exons should show correlated levels of ASE as well as concordant class of genetic divergence (*cis, trans* etc.) when compared within the same F1 individual. It is worth to note that the divergence class also depend on expression levels assigned to the SNP position in parents, which we ignored for simplicity in this analysis.

SNPs assigned to same exons and showing identical type of genetic divergence (concordant SNPs) tended to have strongly correlated ASE levels (Figure S17a). SNPs assigned to same exons but showing different classes of genetic divergence (discordant SNPs) were almost 50% rarer compared to concordant SNPs and, as expected, showed lower level of correlation in ASE and overall smaller allelic differences (Figure S17b). Concordant and discordant SNPs within exons showed overall similar distribution among divergence classes, indicating that discordant calls influenced all classes equally (not shown).

SNPs assigned to the same transcript but different exons had a correlated level of ASE in cases where the divergence class assigned to both exons were the same (Figure S17c). Cases where the exons showed different divergence class showed a lower correlation in ASE (S17d). These results are consistent with previous studies demonstrating variable levels of ASE along genes and between exons (Skelly, Johansson, Madeoy, Wakefield, & Akey, 2011). Different exons of the same transcript that show different levels of ASE suggest that ASE effects are specific to single isoforms rather than all isoforms assigned to the same transcript. Through our RNA-seq analysis we identified over 162000 known and new isoforms distributed to 29296 transcribed loci (on average over 5 isoforms per transcribed locus).

In our final analysis, we opted to classify each transcript into one divergence class, based on the SNP that showed the highest statistical support (see Methods). The justification for this choice was based on the following considerations:

1. Individual SNPs assigned to the same exon tended to show similar ASE levels and divergence types, indicating that dissection of divergence architecture was generally robust to different SNPs within exons. Discordance in divergence types for SNPs assigned to the same exon influenced all divergence types equally and therefore is not expected to bias the results.
2. SNPs assigned to the same transcript but different exons showed different divergence types in roughly half of the cases, and the levels of ASE on the SNP loci were less correlated. This suggests that different exons may experience varying levels of ASE, likely because of alternative isoform expression, as has been demonstrated before (Skelly et al., 2011). Visual inspection of expression tracks in candidate genes for variable ASE identified multiple instances of putative alternative isoform expression (example in Fig S18). Any procedure that would not distinguish different exons, for example averaging expression levels across SNPs, would therefore suffer from low sensitivity as loci not showing ASE would cancel the signal from loci showing ASE.
3. We discarded the option of averaging ASE levels for SNPs assigned to the same exon because different transcripts and ecotype-pairs showed markedly different densities of SNPs. Averaging would therefore influence transcripts and ecotype-pairs unequally.

## References

Bell, M. A., & Foster, S. A. (1994). The evolutionary biology of the threespine stickleback. Oxford science publications (pp. xii–571 p.). Oxford; New York: Oxford University Press.

Brawand, D., Wagner, C. E., Li, Y. I., Malinsky, M., Keller, I., Fan, S., et al. (2014). The genomic substrate for adaptive radiation in African cichlid fish. Nature, 513(7518), 375–381. http://doi.org/10.1038/nature13726

Buil, A., Brown, A. A., Lappalainen, T., Viñuela, A., Davies, M. N., Zheng, H.-F., et al. (2015). Gene-gene and gene-environment interactions detected by transcriptome sequence analysis in twins. Nature Genetics, 47(1), 88–91. http://doi.org/10.1038/ng.3162

Carroll, S. B. (2008). Evo-devo and an expanding evolutionary synthesis: A genetic theory of morphological evolution. Cell, 134(1), 25–36. http://doi.org/10.1016/j.cell.2008.06.030

Chan, Y. F., Marks, M. E., Jones, F. C., Villarreal, G., Shapiro, M. D., Brady, S. D., et al. (2010). Adaptive evolution of pelvic reduction in sticklebacks by recurrent deletion of a Pitx1 enhancer. Science, 327(5963), 302–305.

Cleves, P. A., Ellis, N. A., Jimenez, M. T., Nunez, S. M., Schluter, D., Kingsley, D. M., & Miller, C. T. (2014). Evolved tooth gain in sticklebacks is associated with a cis-regulatory allele of Bmp6. Proceedings of the National Academy of Sciences, 111(38), 13912–13917.

Colosimo, P. F., Hosemann, K. E., Balabhadra, S., Villarreal, G. J., Dickson, M., Grimwood, J., et al. (2005). Widespread parallel evolution in sticklebacks by repeated fixation of Ectodysplasin alleles. Science, 307(5717), 1928–1933. http://doi.org/10.1126/science.1107239

Coolon, J. D., McManus, C. J., Stevenson, K. R., Graveley, B. R., & Wittkopp, P. J. (2014). Tempo and mode of regulatory evolution in Drosophila, 24(5), 797–808.

Cooper, T. F., Rozen, D. E., & Lenski, R. E. (2003). Parallel changes in gene expression after 20,000 generations of evolution in Escherichia coli. Proceedings of the National Academy of Sciences, 100(3), 1072–1077. http://doi.org/10.1073/pnas.0334340100

Danecek, P., Auton, A., Abecasis, G., Albers, C. A., Banks, E., DePristo, M. A., et al. (2011). The variant call format and VCFtools. Bioinformatics, 27(15), 2156–2158. http://doi.org/10.1093/bioinformatics/btr330

Dobin, A., Davis, C. A., Schlesinger, F., Drenkow, J., Zaleski, C., Jha, S., et al. (2013). STAR: ultrafast universal RNA-seq aligner. Bioinformatics, 29(1), 15–21. http://doi.org/10.1093/bioinformatics/bts635

Eden, E., Navon, R., Steinfeld, I., Lipson, D., & Yakhini, Z. (2009). GOrilla: a tool for discovery and visualization of enriched GO terms in ranked gene lists. BMC Bioinformatics, 10(1). http://doi.org/10.1186/1471-2105-10-48

Emerson, J. J., Hsieh, L.-C., Sung, H.-M., Wang, T.-Y., Huang, C.-J., Lu, H. H.-S., et al. (2010). Natural selection on cis and trans regulation in yeasts, Genome Research 20(6), 826–836.

Evans, D. H., Piermarini, P. M., & Choe, K. P. (2005). The multifunctional fish gill: dominant site of gas exchange, osmoregulation, acid-base regulation, and excretion of nitrogenous waste. Physiological Reviews, 85(1), 97–177. http://doi.org/10.1152/physrev.00050.2003

Fraser, H. B., Moses, A. M., & Schadt, E. E. (2010). Evidence for widespread adaptive evolution of gene expression in budding yeast. Proceedings of the National Academy of Sciences, 107(7), 2977–2982.

Gibbons, T. C., Metzger, D. C. H., Healy, T. M., & Schulte, P. M. (2017). Gene expression plasticity in response to salinity acclimation in threespine stickleback ecotypes from different salinity habitats. Molecular Ecology, 34(10), 2711–2725. http://doi.org/10.1111/mec.14065

Gibson, G., Riley-Berger, R., Harshman, L., Kopp, A., Vacha, S., Nuzhdin, S., & Wayne, M. (2004). Extensive sex-specific nonadditivity of gene expression in Drosophila melanogaster. Genetics, 167(4), 1791–1799.

Grossman, S. R., Andersen, K. G., Shlyakhter, I., Tabrizi, S., Winnicki, S., Yen, A., et al. (2013). Identifying Recent Adaptations in Large-Scale Genomic Data. Cell, 152(4), 703–713. http://doi.org/10.1016/j.cell.2013.01.035

Gunter, H. M., Schneider, R. F., Karner, I., Sturmbauer, C., & Meyer, A. (2017). Molecular investigation of genetic assimilation during the rapid adaptive radiations of East African cichlid fishes. Molecular Ecology, 5(3), 457. http://doi.org/10.1111/mec.14405

Haas, B. J., Papanicolaou, A., Yassour, M., Grabherr, M., Blood, P. D., Bowden, J., et al. (2013). De novo transcript sequence reconstruction from RNA-seq using the Trinity platform for reference generation and analysis. Nature Protocols, 8(8), 1494–1512. http://doi.org/10.1038/nprot.2013.084

Haldane, J. B. S. (1927). A mathematical theory of natural and artificial selection, Part V: Selection and mutation. Mathematical Proceedings of the Cambridge Philosophical Society, 23, 838–844.

Hart, J. C., Ellis, N. A., Eisen, M. B., & Miller, C. T. (2018). Convergent evolution of gene expression in two high-toothed stickleback populations. PLoS Genetics, 14(6), e1007443. http://doi.org/10.1371/journal.pgen.1007443

Hartl, D. L., & Clark, A. G. (1997). Principles of population genetics (Vol. 116). Sinauer associates Sunderland.

Hwang, P.-P., Lee, T.-H., & Lin, L.-Y. (2011). Ion regulation in fish gills: recent progress in the cellular and molecular mechanisms. AJP: Regulatory, Integrative and Comparative Physiology, 301(1), R28–R47. http://doi.org/10.1152/ajpregu.00047.2011

Ihaka, R., & Gentleman, R. (1996). R: A language for data analysis and graphics. J Comput Graph Stat, 5(3), 299–314.

Jones, F. C., Grabherr, M. G., Chan, Y. F., Russell, P., Mauceli, E., Johnson, J., et al. (2012). The genomic basis of adaptive evolution in threespine sticklebacks. Nature, 484(7392), 55–61. http://doi.org/10.1038/nature10944

Kingsley, D. M., Zhu, B., Osoegawa, K., de Jong, P. J., Schein, J., Marra, M., et al. (2004). New genomic tools for molecular studies of evolutionary change in threespine sticklebacks. Behaviour, 141(11), 1331–1344. http://doi.org/10.1163/1568539042948150

Landry, C. R., Hartl, D. L., & Ranz, J. M. (2007a). Genome clashes in hybrids: insights from gene expression. Heredity, 99(5), 483–493. http://doi.org/10.1038/sj.hdy.6801045

Landry, C. R., Lemos, B., Rifkin, S. A., Dickinson, W. J., & Hartl, D. L. (2007b). Genetic properties influencing the evolvability of gene expression. Science, 317(5834), 118–121.

Landry, C. R., Wittkopp, P. J., Taubes, C. H., Ranz, J. M., Clark, A. G., & Hartl, D. L. (2005). Compensatory cis-trans evolution and the dysregulation of gene expression in interspecific hybrids of Drosophila. Genetics, 171(4), 1813–1822.

Lemos, B., Araripe, L. O., Fontanillas, P., & Hartl, D. L. (2008). Dominance and the evolutionary accumulation of cis-and trans-effects on gene expression. Proceedings of the National Academy of Sciences, 105(38), 14471–14476.

Li, H. (2013). Aligning sequence reads, clone sequences and assembly contigs with BWA-MEM. arXiv:1303.3997

Love, M. I., Huber, W., & Anders, S. (2014). Moderated estimation of fold change and dispersion for RNA-seq data with DESeq2. Genome Biology, 15(12), 550. http://doi.org/10.1186/s13059-014-0550-8

McCairns, R. J. S., & Bernatchez, L. (2010). Adaptive divergence between freshwater and marine sticklebacks: insights into the role of phenotypic plasticity from an integrated analysis of candidate gene expression. Evolution, 64(4), 1029–1047. http://doi.org/10.1111/j.1558-5646.2009.00886.x

McKenna, A., Hanna, M., Banks, E., Sivachenko, A., Cibulskis, K., Kernytsky, A., et al. (2010). The Genome Analysis Toolkit: a MapReduce framework for analyzing next-generation DNA sequencing data. Genome Research, 20(9), 1297–1303. http://doi.org/10.1101/gr.107524.110

McManus, C. J., Coolon, J. D., Duff, M. O., Eipper-Mains, J., Graveley, B. R., & Wittkopp, P. J. (2010). Regulatory divergence in Drosophila revealed by mRNA-seq. Genome Research, 20(6), 816–825.

Nielsen, R., Williamson, S., Kim, Y., Hubisz, M. J., Clark, A. G., & Bustamante, C. (2005). Genomic scans for selective sweeps using SNP data. Genome Research, 15(11), 1566–1575. http://doi.org/10.1101/gr.4252305

O’Brown, N. M., Summers, B. R., Jones, F. C., Brady, S. D., & Kingsley, D. M. (2015). A recurrent regulatory change underlying altered expression and Wnt response of the stickleback armor plates gene EDA. Elife, 4. http://doi.org/10.7554/eLife.05290

Orr, H. A. (1998). Testing natural selection vs. genetic drift in phenotypic evolution using quantitative trait locus data. Genetics, 149(4), 2099–2104.

Prud’homme, B., Gompel, N., & Carroll, S. B. (2007). Emerging principles of regulatory evolution. Proceedings of the National Academy of Sciences, 104(Suppl 1), 8605–8612.

Shapiro, M. D., Marks, M. E., Peichel, C. L., Blackman, B. K., Nereng, K. S., Jónsson, B., et al. (2004). Genetic and developmental basis of evolutionary pelvic reduction in threespine sticklebacks. Nature, 428(6984), 717–723. http://doi.org/10.1038/nature02415

Skelly, D. A., Johansson, M., Madeoy, J., Wakefield, J., & Akey, J. M. (2011). A powerful and flexible statistical framework for testing hypotheses of allele-specific gene expression from RNA-seq data. Genome Research, 21(10), 1728–1737. http://doi.org/10.1101/gr.119784.110

Stern, D. L., & Orgogozo, V. (2009). Is genetic evolution predictable? Science, 323(5915), 746–751. http://doi.org/10.1126/science.1158997

Storey, J. D. (2002). A direct approach to false discovery rates. Journal of the Royal Statistical Society: Series B (Statistical Methodology), 64(3), 479–498. http://doi.org/10.1111/1467-9868.00346

Trapnell, C., Roberts, A., Goff, L., Pertea, G., Kim, D., Kelley, D. R., et al. (2012). Differential gene and transcript expression analysis of RNA-seq experiments with TopHat and Cufflinks. Nature Protocols, 7(3), 562–578. http://doi.org/10.1038/nprot.2012.016

Velotta, J. P., Wegrzyn, J. L., Ginzburg, S., Kang, L., Czesny, S., O’Neill, R. J., et al. (2016). Transcriptomic imprints of adaptation to fresh water: parallel evolution of osmoregulatory gene expression in the Alewife. Molecular Ecology, 26(3), 831–848. http://doi.org/10.1111/mec.13983

Verta, J.-P., Landry, C. R., & Mackay, J. (2016). Dissection of expression-quantitative trait locus and allele specificity using a haploid/diploid plant system – insights into compensatory evolution of transcriptional regulation within populations. The New Phytologist, 211(1), 159–171. http://doi.org/10.1111/nph.13888

Waddington, C. H. (1942). Canalization of development and the inheritance of acquired characters. Nature, 150(3811), 563–565. http://doi.org/10.1038/150563a0

Weir, B. S., & Cockerham, C. C. (1984). Estimating F-statistics for the analysis of population structure. Evolution, 38(6), 1358–1370. http://doi.org/10.2307/2408641

Whitehead, A. (2010). The evolutionary radiation of diverse osmotolerant physiologies in killifish (Fundulus sp.). Evolution, 64(7), 2070–2085. http://doi.org/10.1111/j.1558-5646.2010.00957.x

Wittkopp, P. J., Haerum, B. K., & Clark, A. G. (2004). Evolutionary changes in cis and trans gene regulation. Nature, 430(6995), 85–88.

Wittkopp, P. J., Haerum, B. K., & Clark, A. G. (2008). Regulatory changes underlying expression differences within and between Drosophila species. Nature Genetics, 40(3), 346–350. http://doi.org/10.1038/ng.77

