## Supplementary material for "Predominance of *cis*-regulatory changes in parallel expression divergence of sticklebacks"

### Supplemental Figures

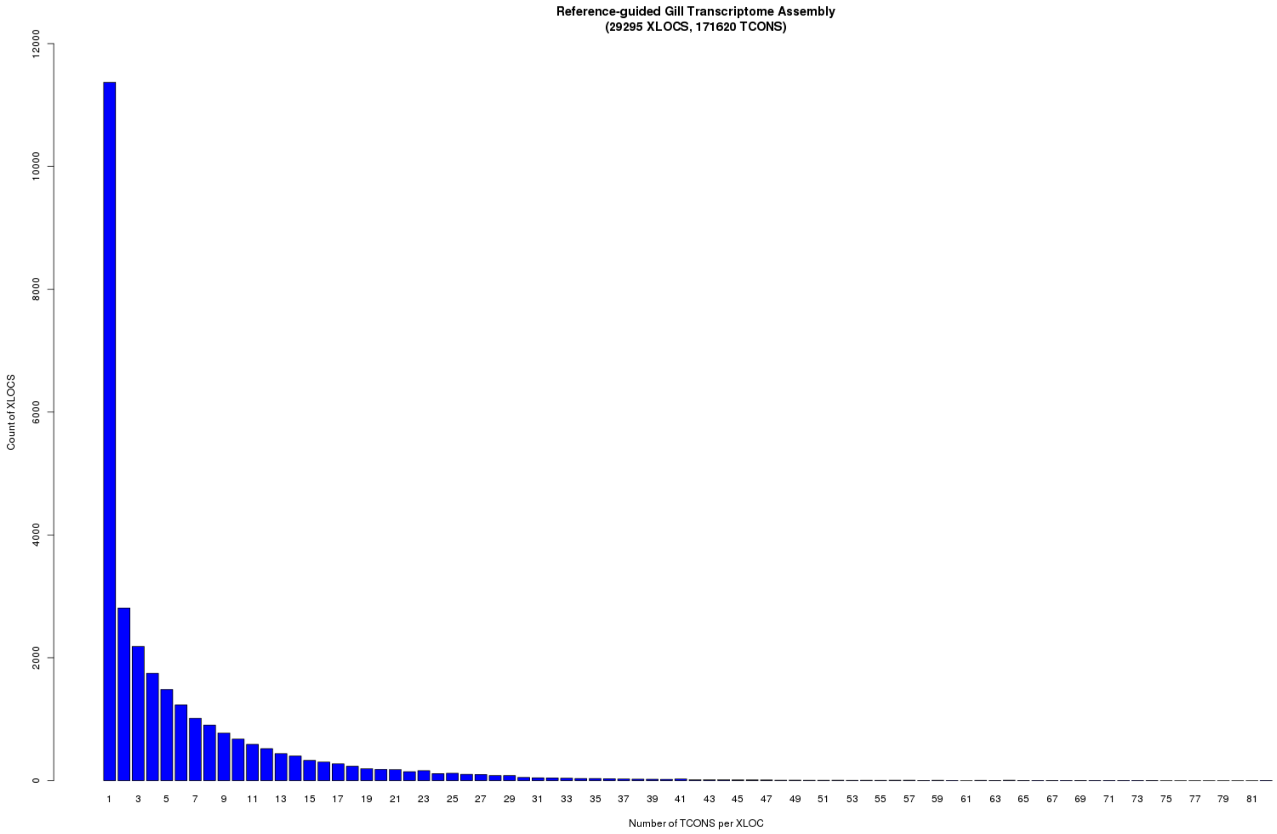


Fig S1. The stickleback gill transcriptome has a highly skewed distribution of isoform variation (number of "TCONS") per expressed locus ("XLOC"). While some loci have >50 isoforms per locus, the majority of loci have 3 (median) or less transcripts per locus.

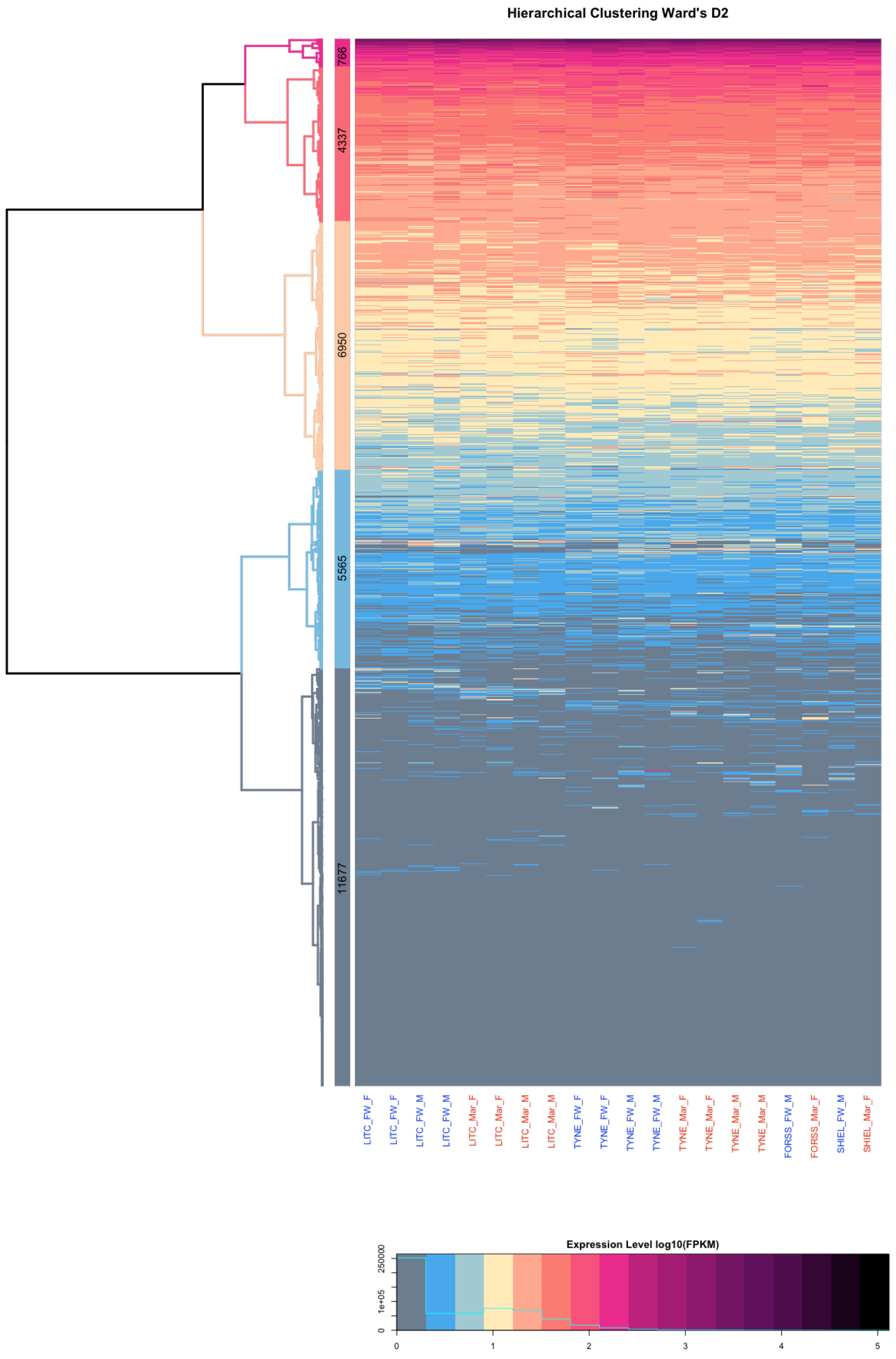


Fig S2. Hierarchical clustering of stickleback gill transcriptome by expression level in 20 marine and freshwater fish reared under standard laboratory conditions reveals five major groups of loci based on their their average expression level. The most highly expressed group of genes showing strong enrichment for biological processes with the respiratory function of the gill including mitochondrial respiration, ATP synthesis coupled proton transport and cytoplasmic translation (Supplementary note).

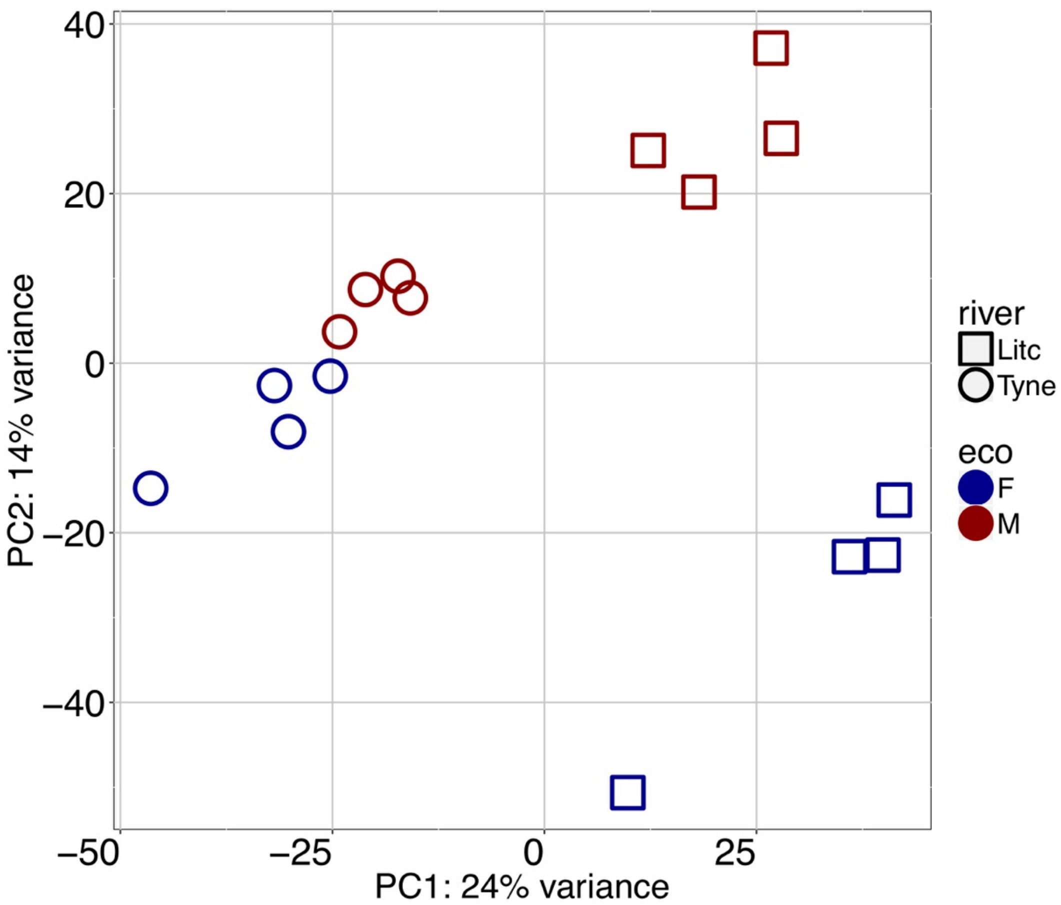


Fig S3. Principal component analysis of gill transcriptomes of Little Campbell and Tyne ecotypes showing principal components 1 and 2 separating individuals by geography (PC1) and ecotype (PC2) respectively.

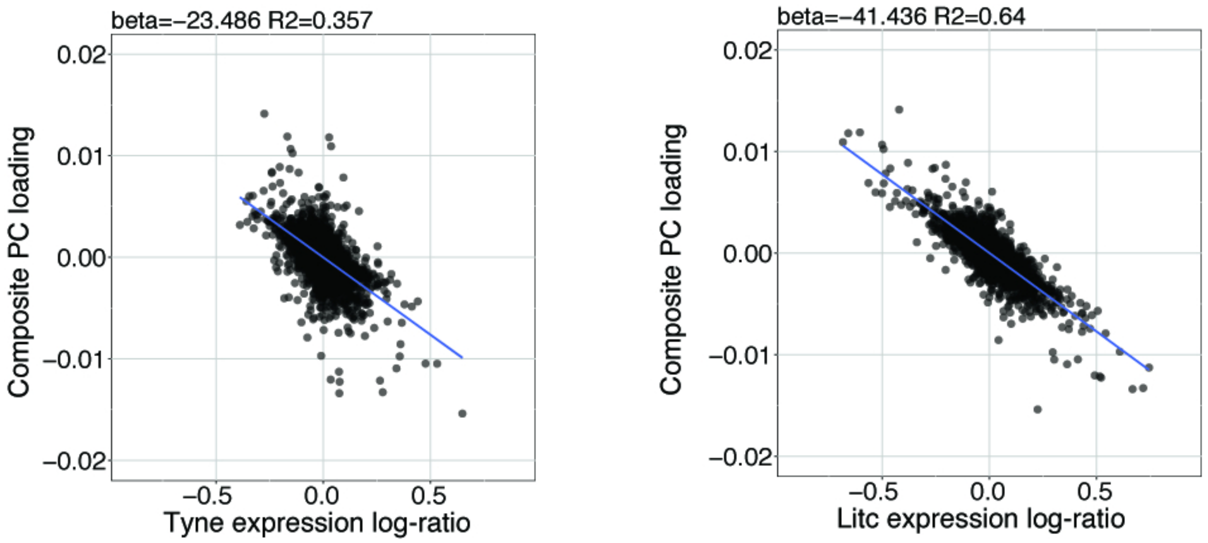


Fig S4. Parallel marine-freshwater divergence in gene expression quantified using the Composite PC loadings (y-axis, see also Fig 1c) is highly correlated with river-specific marine-freshwater divergence in gene expression River Tyne, Scotland (left) and Little Campbell River, Canada (right) quantified as the log of marine/freshwater expression values. Each point represents a transcript. Negative values on x- and y-axes represent stronger expression in freshwater fish and positive values represent stronger expression in marine fish.

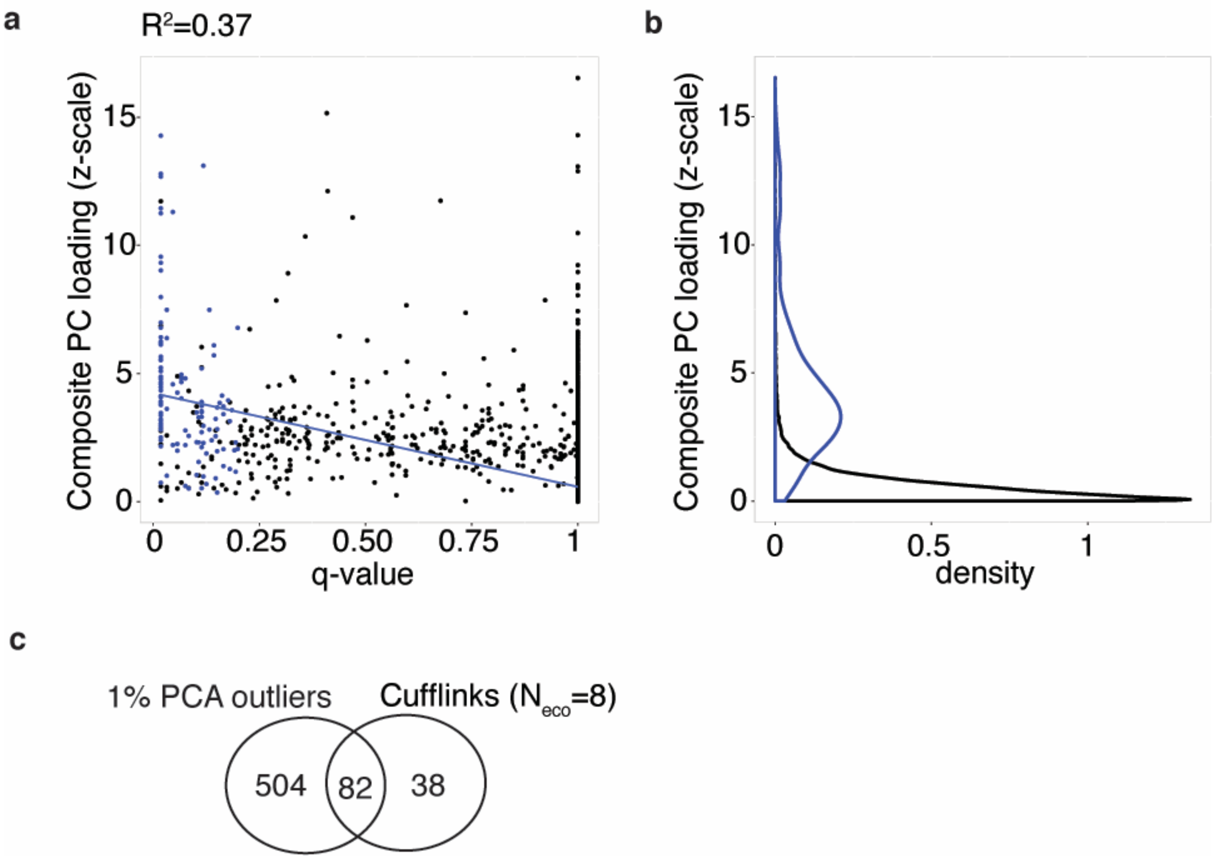


Fig S5. (ab) Parallel freshwater-marine differentially expressed transcripts identified using parametric tests (FDR 20%, blue points and density estimation) versus parallel differentially expressed transcripts identified from composite PC loadings. Variation in composite PC loading explains 37% of variation in q-value (blue line). (c) Sharing of parallel freshwater-marine differentially expressed transcripts versus transcripts that have the strongest contribution to sample separation on composite PC ("PC outliers", i.e. composite PC loading in the lowest or highest 1% of loading distribution and correlated freshwater-marine mean expression difference in Little Campbell and Tyne).

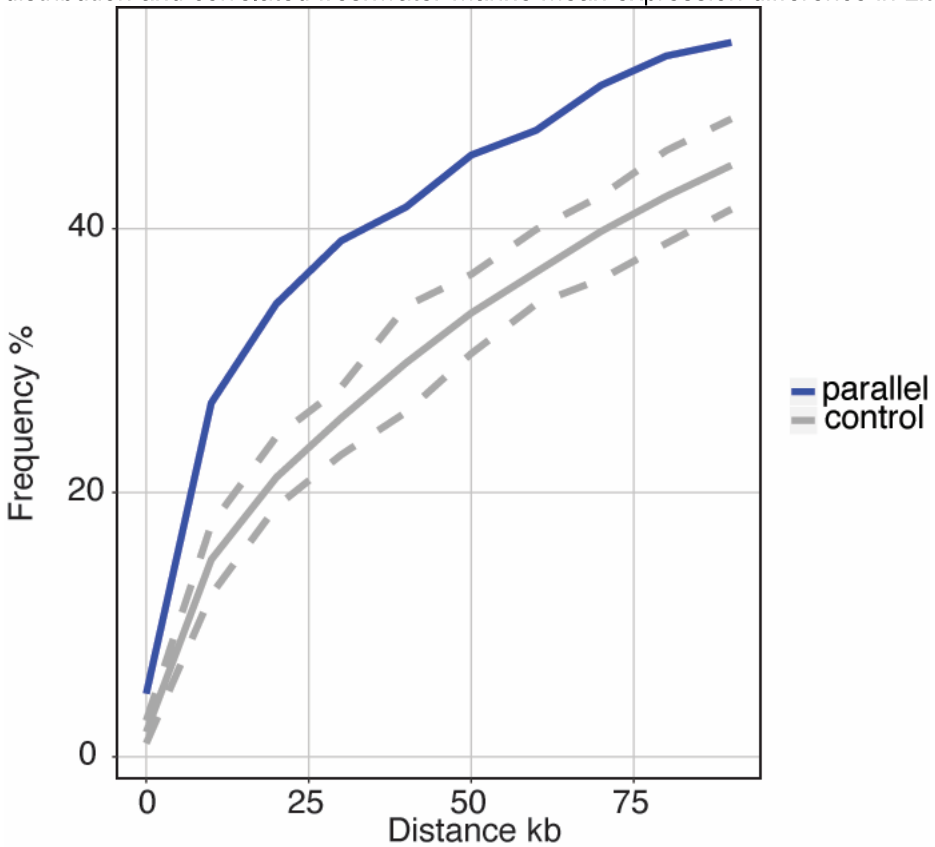


Fig S6. Transcripts with parallel expression divergence are in close spatial proximity to regions of the genome that show elevated parallel genetic divergence (CSS outlier loci) among marine and freshwater fish from the River Tyne and Little Campbell that were genome sequenced in this study.

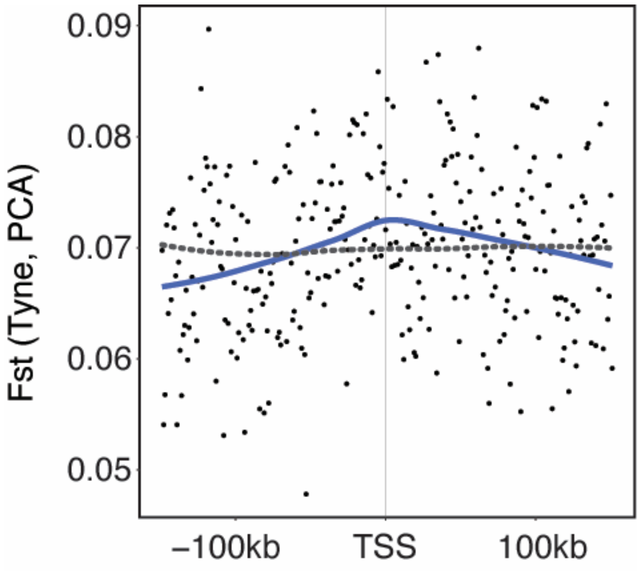

Fig S7. Genetic divergence ($F_{ST}$) calculated in 1kb windows around the 586 loci with parallel divergent expression (black points with blue line representing mean) relative to randomly sampled loci (grey dashed line) in Tyne populations.

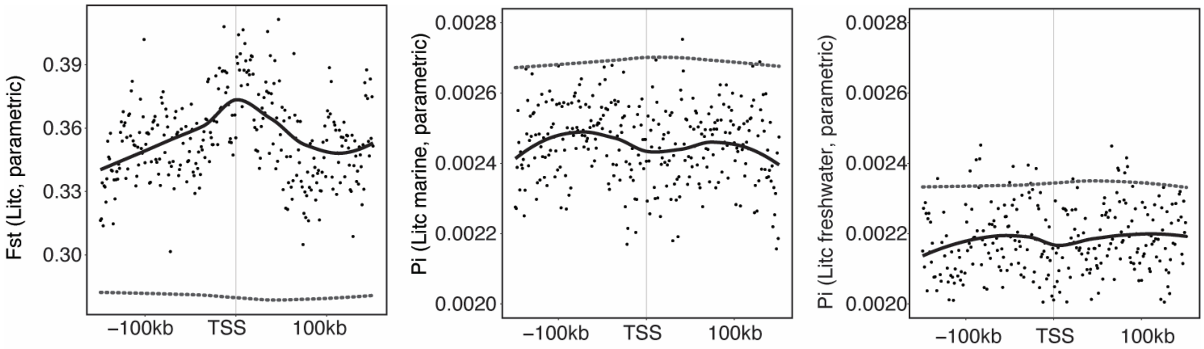


Fig S8. Genetic divergence (Fst) and nucleotide diversity (Pi) associated with loci showing parallel expression divergence defined based on a parametric test performed in cufflinks (Little Campbell). Black points represent values for 1kb windows, and solid line a loess smoothed mean. Dashed grey lines represent mean $F_{ST}$ for randomly sampled 'control' windows. Results for Tyne were non-significant (not shown).

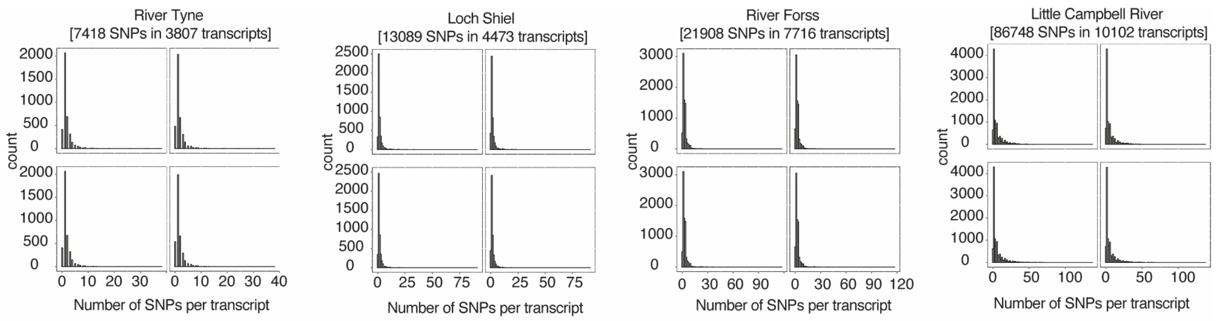


Fig S9. Number of informative (tested) SNPs per transcript per cross. Four panels per cross represent four analyzed F1's. Zero corresponds to positions where SNP position was not informative (expressed) in a given F1, but informative in other F1's of the same family.

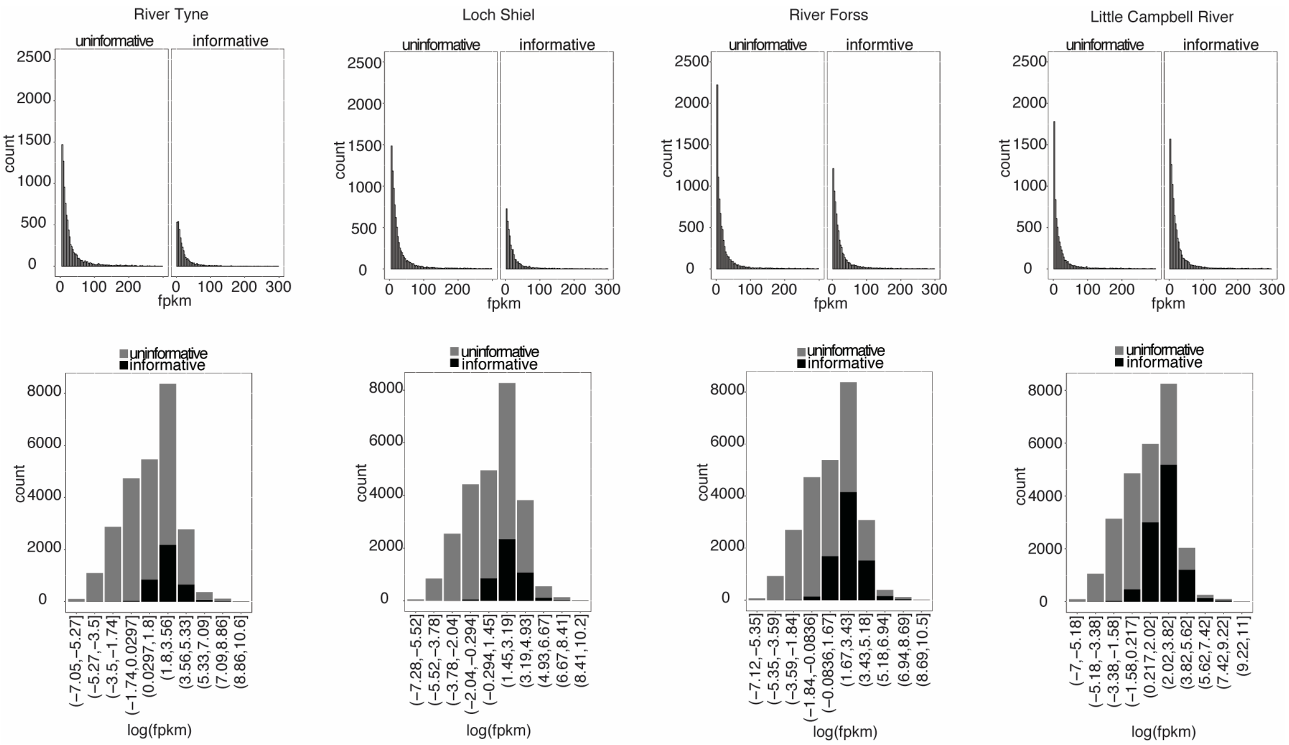


Fig S10. Expression levels of transcripts with informative SNPs. Average FPKM values in transcripts with informative SNPs for allele-specific expression test and those without informative SNPs (uninformative).

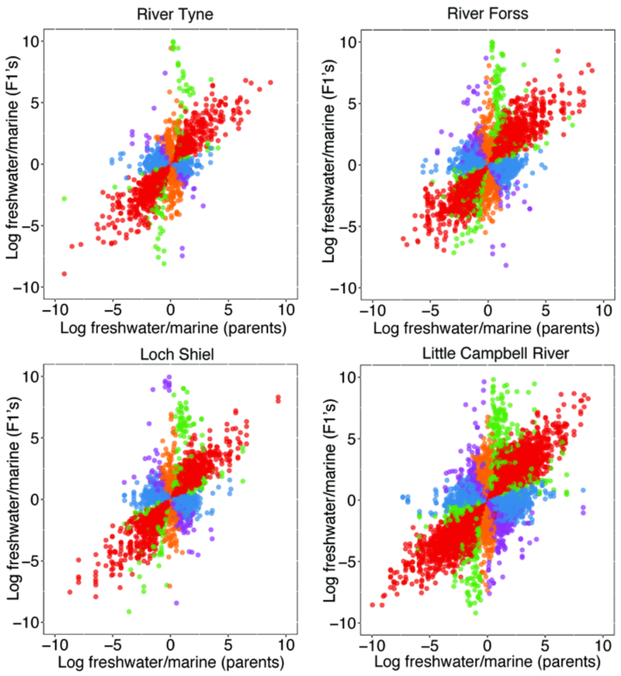


Fig S11. The genetic architecture of expression divergence between ecotype pairs in all investigated ecotype-pairs (as per Fig 3b).

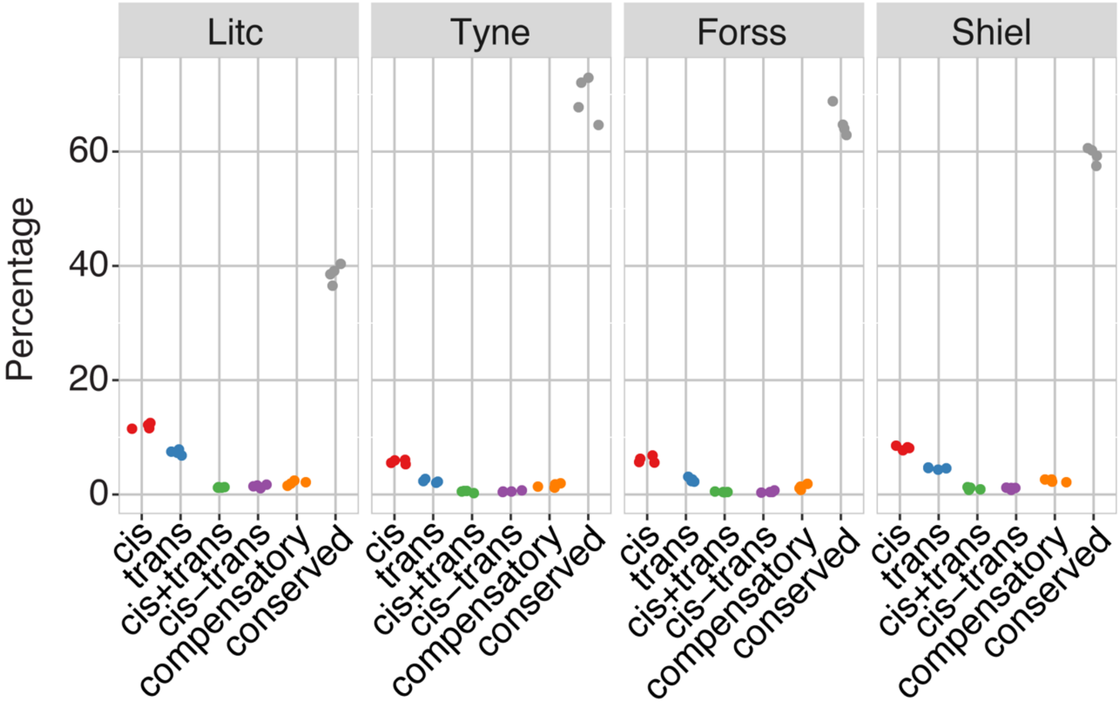


Fig S12. Frequencies of genetic architectures of expression divergence in down-sampled (30M uniquely mapping reads per sample) dataset.

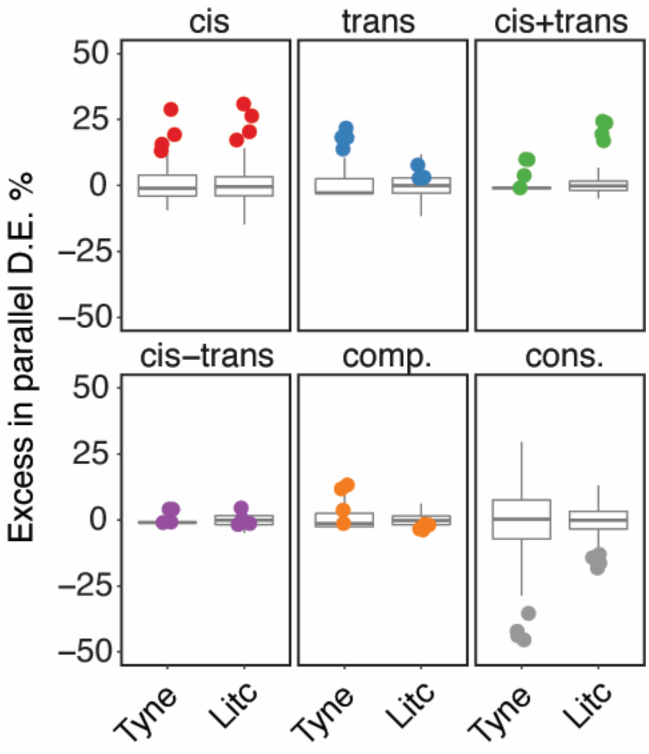

Fig S13. Cis/trans-overrepresentation (as per Fig 4) in parallel evolving genes as defined by parametric test.

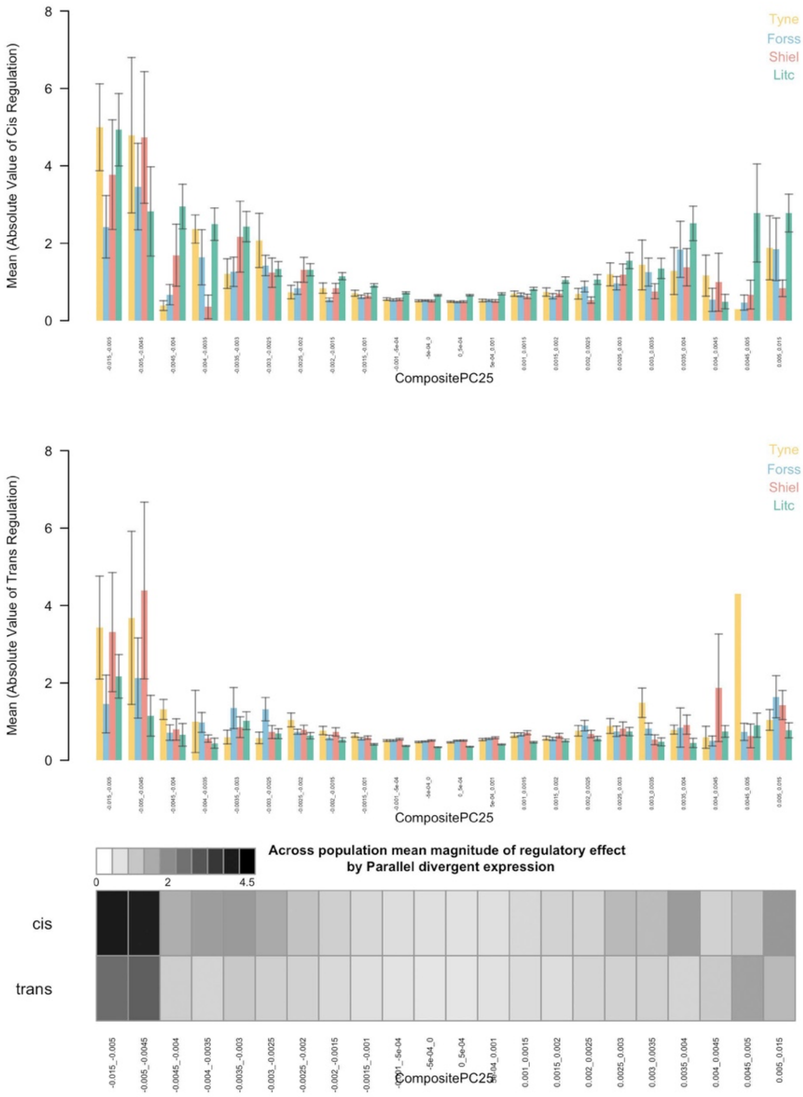


Figure S14. Mean magnitude of cis- (a) and trans- (b) regulation relative to parallel divergence in gene expression measured by composite PC. For each population studied for allele-specific expression, loci were binned according to their loadings on the composite PC describing parallel divergence in gene expression of pure strains. For each bin and each population the mean and standard error of the quantitative degree of cis-regulation and trans-regulation was calculated (see main text). (c) As per Fig5c in main text. For each bin, the mean across populations was calculated and plotted as a heat map with darker grey representing larger magnitudes of cis- and trans- effects.

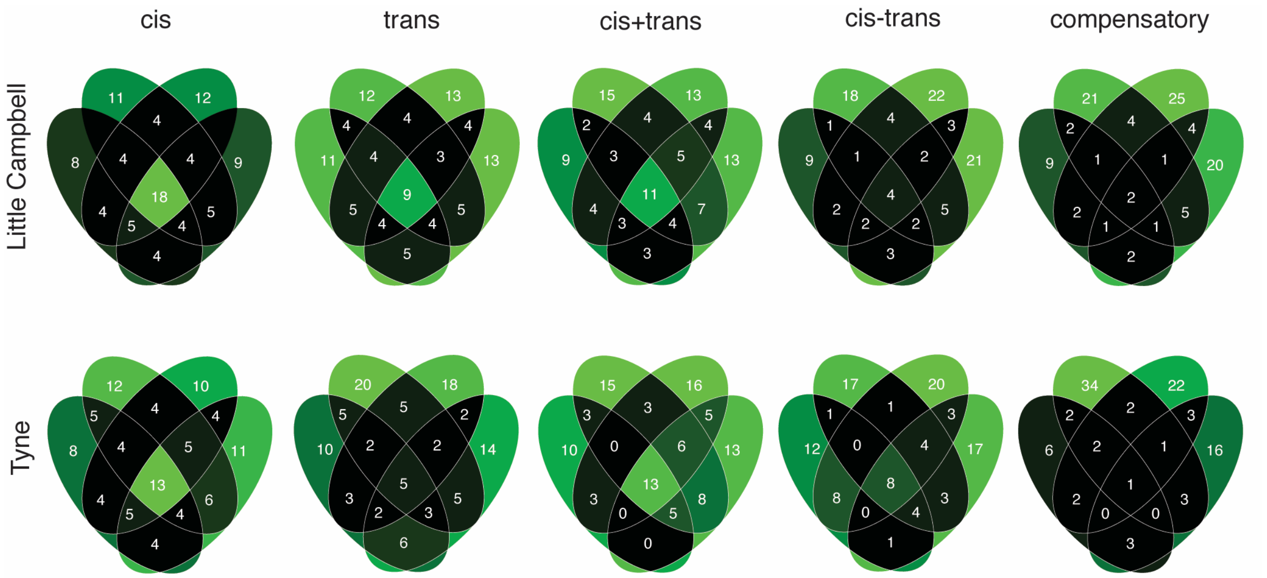


Fig S15. Sharing of regulatory divergence between siblings as a proxy for the level of epistasis associated with different genetic architectures in Little Campbell (see also main text Fig 6b) and Tyne. The type of regulation observed for each gene is compared among siblings (overlapping ovals) with numbers representing percent of loci shared between siblings (rounded to integer). Color scale ranges from black (low) to pale green (high). Cis-regulatory divergence tended to be most stable across genetic backgrounds indicating that cis-acting divergence is least influenced by epistasis.

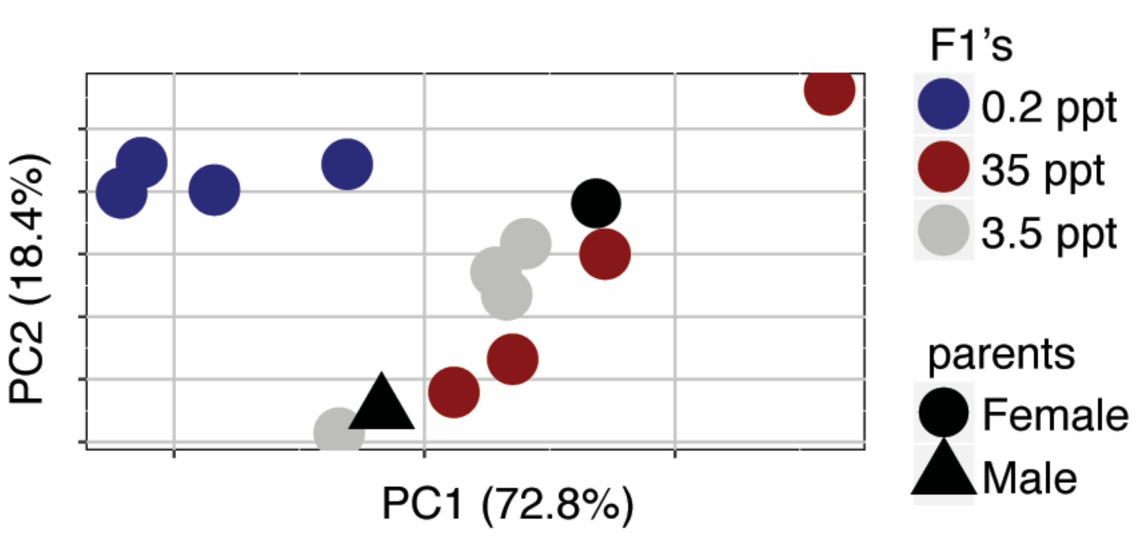


Fig S16. Principal Component Analysis of expression profiles of full siblings from a Little Campbell River marine x freshwater cross acclimated to different salinities.

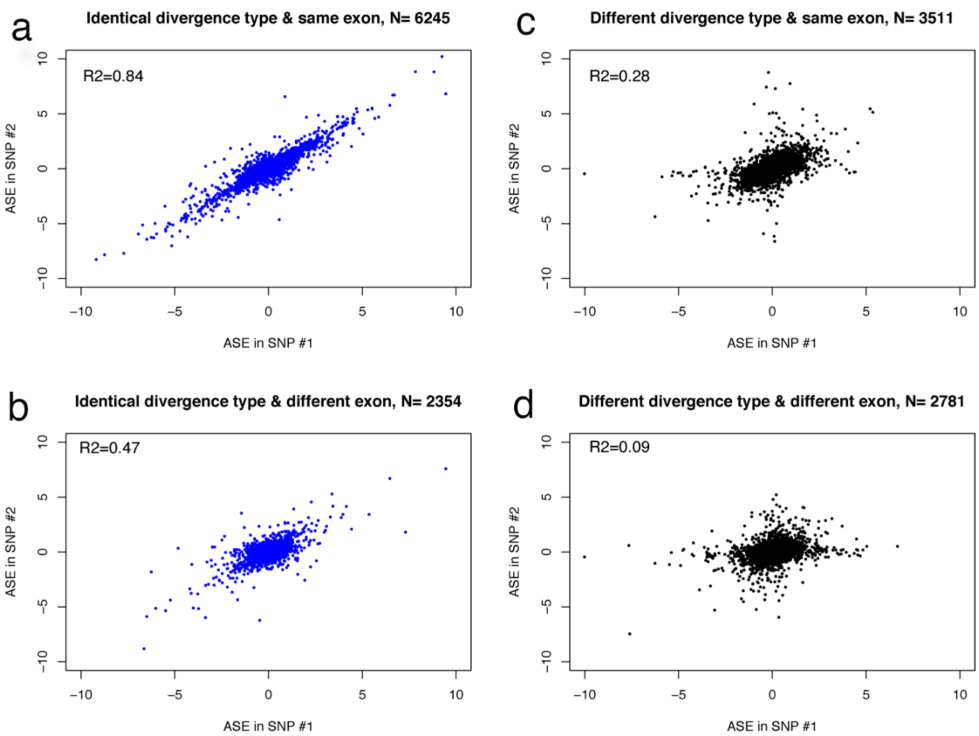


Fig S17. Correlation of ASE (log-fold change of RNA-seq counts of marine over freshwater allele) in different pairs of SNPs was used to perform concordance analysis. (a) SNPs assigned to same exons and showing identical type of genetic divergence (concordant SNPs). (b) SNPs assigned to same exons but showing different classes of genetic divergence (discordant SNPs). (c) SNPs assigned to different exons of the same transcript and showing identical type of genetic divergence. (d) SNPs assigned to different exons of the same transcript and showing different type of genetic divergence.

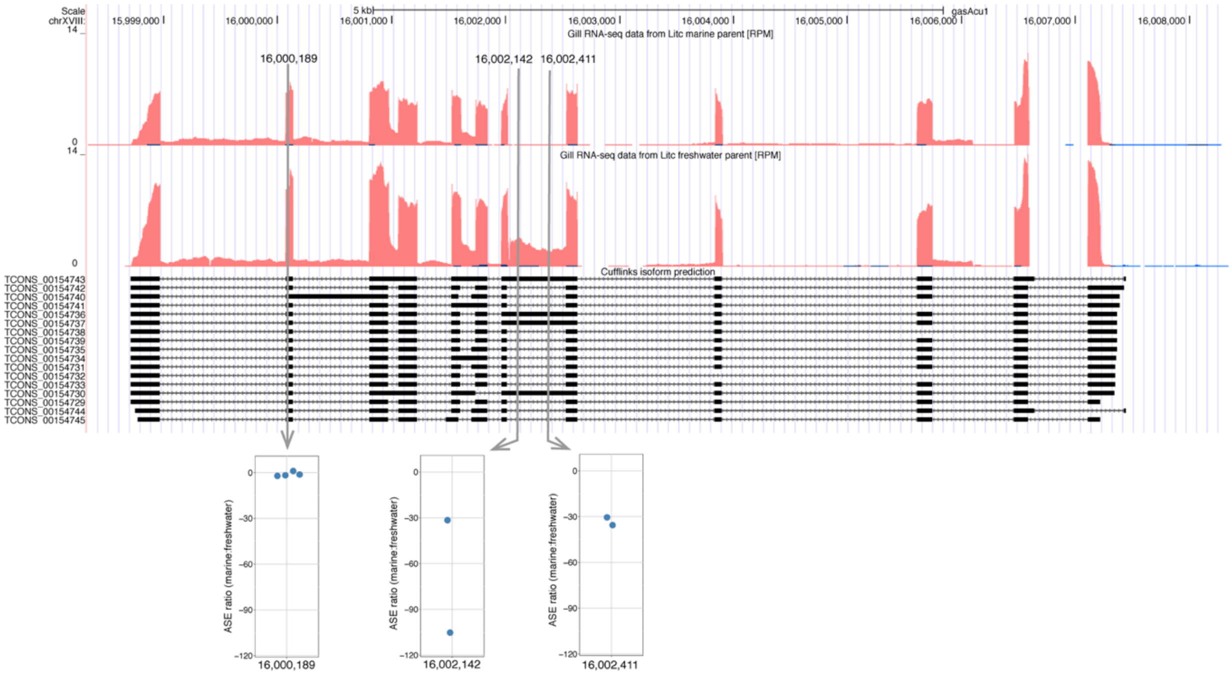


Figure S18. Candidate transcript for alternative isoform expression. This example illustrates that SNPs in different regions of the same transcript can show varying levels of ASE associated with alternative splice forms. Expression level is represented by RPM (Reads [mapping to position] Per Million [reads mapped overall]) and corresponds to red track in Little Campbell marine (upper) and freshwater (lower) parents. CuffLinks isoform predictions are represented below expression tracks. The marine and freshwater alleles are distinguished by three SNPs mapping to two exons. SNP at position chrXVIII:16000189 shows similar expression level of both alleles and hence no ASE. SNPs at positions 16002142 and 16002411 map to an alternatively spliced region of the transcript where expression is observed in freshwater allele but not in marine allele, and hence the SNPs show ASE towards freshwater.
