## Supplementary material for "Predominance of *cis*-regulatory changes in parallel expression divergence of sticklebacks"

### Supplemental Tables

| Table S1. Sampling locations | | |
|---|---|---|
| Sampling locations | Freshwater sampling coordinates | Marine sampling coordinates |
| River Tyne | 55°56'34.8"N 2°47'6"W | 56°0'22.63"N 2°36'47.41"W |
| River Forss | 58°34'22.949"N 3°38'27.916"W | 58°36'32.335"N 3°40'42.404"W |
| River Shiel | 56°44'52.8"N 5°41'52.8"W | 56°46'45.99"N 5°49'44.47"W |
| Little Campbell River | 49°0'43.65"N 122°37'30.44"W | 49°0'58.03"N 122°46'46.37"W |

| Table S2. RNA-seq library sequencing and yield. | | | | | |
|---|---|---|---|---|---|
| **Sample ID** | **Sampling location** | **Ecotype** | **Sequencing run** | **Sequencing lane** | **Mapped reads *** |
| c363_P_FC18_M | Little Campbell River | Freshwater (parent) | 48 | 1 | 41570029 |
| c363_P_FC08_F | Little Campbell River | Marine (parent) | 48 | 1 | 77071816 |
| c358_P_FC12_F | Little Campbell River | Freshwater | 48 | 2 | 51196050 |
| c358_P_FC09_M | Little Campbell River | Marine | 48 | 2 | 62721395 |
| c357_P_FC14_F | Little Campbell River | Freshwater | 48 | 2 | 51219036 |
| c357_P_FC06_M | Little Campbell River | Marine | 48 | 2 | 49172261 |
| c353_P_FC15_M | Little Campbell River | Freshwater | 48 | 2 | 62929997 |
| c353_P_FC05_F | Little Campbell River | Marine | 48 | 2 | 40600091 |
| c209_P_422_M | River Tyne | Freshwater | 39 | 7 | 57545801 |
| c209_P_341_FM | River Tyne | Marine | 39 | 7 | 65354927 |
| c208_P_531_M | River Tyne | Marine | 39 | 7 | 51827643 |
| c208_P_321_FM | River Tyne | Freshwater | 39 | 7 | 59257432 |
| c172_P_533_M | River Tyne | Freshwater (parent) | 14 | 8 | 37475539 |
| c172_P_533_M | River Tyne | Freshwater (parent) | 14 | 3 | 37561905 |
| c172_P_532_F | River Tyne | Marine (parent) | 48 | 2 | 32355769 |
| c172_P_532_F | River Tyne | Marine (parent) | 14 | 8 | 29421104 |
| c172_P_532_F | River Tyne | Marine (parent) | 14 | 3 | 30491397 |
| c169_P_432_FM | River Tyne | Freshwater | 39 | 7 | 46449279 |
| c169_P_342_M | River Tyne | Marine | 39 | 7 | 60308422 |
| c214_P_524_FM | River Shiel | Marine (parent) | 39 | 6 | 63867976 |
| c214_P_512_M | River Shiel | Freshwater (parent) | 39 | 6 | 51993114 |
| c212_P_551_M | River Forss | Freshwater (parent) | 39 | 5 | 78491454 |
| c212_P_454_FM | River Forss | Marine (parent) | 39 | 5 | 53915757 |
| c363_F1_1_M | Little Campbell River | F1 | 48 | 1 | 51918812 |
| c363_F1_1_F | Little Campbell River | F1 | 48 | 1 | 75548850 |
| c363_F1_2_M | Little Campbell River | F1 | 48 | 1 | 56909054 |
| c363_F1_2_F | Little Campbell River | F1 | 48 | 1 | 49711377 |
| c172_F1_04_M | River Tyne | F1 | 14 | 8 | 43447943 |
| c172_F1_04_F | River Tyne | F1 | 14 | 8 | 30068470 |
| c172_F1_20_M | River Tyne | F1 | 14 | 3 | 26736357 |
| c172_F1_20_F | River Tyne | F1 | 14 | 3 | 42657169 |
| c214_F1_2_M | River Shiel | F1 | 39 | 6 | 67202528 |
| c214_F1_2_FM | River Shiel | F1 | 39 | 6 | 51160598 |
| c214_F1_1_M | River Shiel | F1 | 39 | 6 | 58378310 |
| c214_F1_1_FM | River Shiel | F1 | 39 | 6 | 62005808 |
| c212_F1_2_M | River Forss | F1 | 39 | 5 | 52427751 |
| c212_F1_2_FM | River Forss | F1 | 39 | 5 | 62315657 |
| c212_F1_1_M | River Forss | F1 | 39 | 5 | 56085728 |
| c212_F1_1_FM | River Forss | F1 | 39 | 5 | 53416036 |

* uniquely mapped reads

**Table S3. Gene Ontology term enrichment.** Available as a supplementary xls file. Two different Gene Ontology analyses were performed using parallel divergent gene expression outliers defined through parametric analysis (CuffDiff) and composite PC analysis (compPC). Gene ontologies with significance lower than $1x10^{-5}$ shown.

| Analysis | GO Term | Description | P-value | FDR q-value | Enrich-ment | N | B | n | b |
|---|---|---|---|---|---|---|---|---|---|
| CuffDiff | GO:0051049 | regulation of transport | 2,76E-10 | 3,87E-06 | 1,59 | 1131 5 | 1295 | 1027 | 187 |
| CuffDiff | GO:0043269 | regulation of ion transport | 8,87E-10 | 6,22E-06 | 2,07 | 1131 5 | 395 | 1134 | 82 |
| CuffDiff | GO:0065008 | regulation of biological quality | 1,74E-09 | 8,13E-06 | 1,38 | 1131 5 | 2189 | 1148 | 307 |
| CuffDiff | GO:0007186 | G-protein coupled receptor signaling pathway | 2,19E-09 | 7,69E-06 | 2,01 | 1131 5 | 435 | 1100 | 85 |
| CuffDiff | GO:0006811 | ion transport | 1,45E-08 | 4,06E-05 | 1,72 | 1131 5 | 769 | 1036 | 121 |
| CuffDiff | GO:0034765 | regulation of ion transmembrane transport | 7,23E-08 | 1,69E-04 | 2,34 | 1131 5 | 209 | 1134 | 49 |
| CuffDiff | GO:0034762 | regulation of transmembrane transport | 1,73E-07 | 3,47E-04 | 2,28 | 1131 5 | 214 | 1134 | 49 |
| CuffDiff | GO:0003008 | system process | 2,15E-07 | 3,76E-04 | 1,58 | 1131 5 | 878 | 1134 | 139 |
| CuffDiff | GO:0055082 | cellular chemical homeostasis | 5,37E-07 | 8,37E-04 | 1,83 | 1131 5 | 435 | 1138 | 80 |
| CuffDiff | GO:0048878 | chemical homeostasis | 7,20E-07 | 1,01E-03 | 1,66 | 1131 5 | 640 | 1138 | 107 |
| CuffDiff | GO:0042391 | regulation of membrane potential | 8,20E-07 | 1,05E-03 | 2 | 1131 5 | 301 | 1148 | 61 |
| CuffDiff | GO:0034220 | ion transmembrane transport | 8,62E-07 | 1,01E-03 | 1,87 | 1131 5 | 390 | 1134 | 73 |
| CuffDiff | GO:0023061 | signal release | 8,80E-07 | 9,49E-04 | 2,77 | 1131 5 | 114 | 1148 | 32 |
| CuffDiff | GO:0001505 | regulation of neurotransmitter levels | 1,07E-06 | 1,08E-03 | 2,33 | 1131 5 | 176 | 1131 | 41 |
| CuffDiff | GO:0043270 | positive regulation of ion transport | 1,19E-06 | 1,11E-03 | 2,42 | 1131 5 | 157 | 1134 | 38 |
| CuffDiff | GO:0006941 | striated muscle contraction | 1,27E-06 | 1,11E-03 | 3,55 | 1131 5 | 59 | 1134 | 21 |
| CuffDiff | GO:0060048 | cardiac muscle contraction | 1,59E-06 | 1,31E-03 | 4,54 | 1131 5 | 33 | 1134 | 15 |
| CuffDiff | GO:0019725 | cellular homeostasis | 1,60E-06 | 1,25E-03 | 1,72 | 1131 5 | 513 | 1138 | 89 |
| CuffDiff | GO:0055085 | transmembrane transport | 2,25E-06 | 1,66E-03 | 1,7 | 1131 5 | 591 | 1036 | 92 |
| CuffDiff | GO:0042592 | homeostatic process | 2,45E-06 | 1,72E-03 | 1,56 | 1131 5 | 914 | 981 | 124 |
| CuffDiff | GO:0006936 | muscle contraction | 2,81E-06 | 1,88E-03 | 2,69 | 1131 5 | 115 | 1134 | 31 |
| CuffDiff | GO:0050801 | ion homeostasis | 2,99E-06 | 1,91E-03 | 1,75 | 1131 5 | 467 | 1138 | 82 |
| CuffDiff | GO:0003012 | muscle system process | 3,02E-06 | 1,84E-03 | 2,39 | 1131 5 | 150 | 1134 | 36 |
| CuffDiff | GO:0015672 | monovalent inorganic cation transport | 3,16E-06 | 1,85E-03 | 2,14 | 1131 5 | 222 | 1094 | 46 |
| CuffDiff | GO:0098771 | inorganic ion homeostasis | 3,84E-06 | 2,15E-03 | 1,75 | 1131 5 | 448 | 1138 | 79 |
| CuffDiff | GO:0032879 | regulation of localization | 4,77E-06 | 2,57E-03 | 1,35 | 1131 5 | 1849 | 1037 | 229 |
| CuffDiff | GO:0098655 | cation transmembrane transport | 6,37E-06 | 3,31E-03 | 1,97 | 1131 5 | 274 | 1134 | 54 |

| | | | | | | | | | |
|---|---|---|---|---|---|---|---|---|---|
| CuffDiff | GO:0010959 | regulation of metal ion transport | 9,75E-06 | 4,89E-03 | 2,1 | 1131 5 | 217 | 1094 | 44 |
| CuffDiff | GO:0051480 | regulation of cytosolic calcium ion concentration | 9,78E-06 | 4,73E-03 | 2,16 | 1131 5 | 189 | 1138 | 41 |
| CuffDiff | GO:0007626 | locomotory behavior | 9,85E-06 | 4,60E-03 | 2,2 | 1131 5 | 180 | 1115 | 39 |
| CompPC | GO:0060048 | cardiac muscle contraction | 1,40E-08 | 1,97E-04 | 5,86 | 1131 5 | 33 | 936 | 16 |
| CompPC | GO:1903522 | regulation of blood circulation | 9,70E-08 | 6,80E-04 | 3,03 | 1131 5 | 112 | 1066 | 32 |
| CompPC | GO:0008016 | regulation of heart contraction | 2,36E-07 | 1,10E-03 | 2,99 | 1131 5 | 110 | 1066 | 31 |
| CompPC | GO:0051239 | regulation of multicellular organismal process | 3,57E-07 | 1,25E-03 | 1,43 | 1131 5 | 1446 | 1109 | 203 |
| CompPC | GO:0050801 | ion homeostasis | 5,20E-07 | 1,46E-03 | 2,01 | 1131 5 | 467 | 770 | 64 |
| CompPC | GO:0044057 | regulation of system process | 7,42E-07 | 1,74E-03 | 1,96 | 1131 5 | 340 | 1106 | 65 |
| CompPC | GO:0065008 | regulation of biological quality | 8,76E-07 | 1,76E-03 | 1,34 | 1131 5 | 2189 | 1022 | 265 |
| CompPC | GO:0050727 | regulation of inflammatory response | 1,24E-06 | 2,17E-03 | 2,63 | 1131 5 | 151 | 940 | 33 |
| CompPC | GO:0098771 | inorganic ion homeostasis | 1,39E-06 | 2,17E-03 | 2 | 1131 5 | 448 | 770 | 61 |
| CompPC | GO:0019725 | cellular homeostasis | 1,84E-06 | 2,58E-03 | 1,95 | 1131 5 | 513 | 725 | 64 |
| CompPC | GO:0055080 | cation homeostasis | 2,11E-06 | 2,68E-03 | 2 | 1131 5 | 434 | 770 | 59 |
| CompPC | GO:0006814 | sodium ion transport | 2,88E-06 | 3,37E-03 | 5,99 | 1131 5 | 89 | 276 | 13 |
| CompPC | GO:0042592 | homeostatic process | 3,65E-06 | 3,94E-03 | 1,64 | 1131 5 | 914 | 770 | 102 |
| CompPC | GO:0006941 | striated muscle contraction | 3,99E-06 | 4,00E-03 | 4,1 | 1131 5 | 59 | 795 | 17 |
| CompPC | GO:0006873 | cellular ion homeostasis | 4,45E-06 | 4,16E-03 | 2,06 | 1131 5 | 394 | 725 | 52 |
| CompPC | GO:0048002 | antigen processing and presentation of peptide antigen | 6,21E-06 | 5,44E-03 | 49,63 | 1131 5 | 16 | 57 | 4 |
| CompPC | GO:0006952 | defense response | 6,95E-06 | 5,73E-03 | 1,75 | 1131 5 | 433 | 1135 | 76 |
| CompPC | GO:0006955 | immune response | 9,63E-06 | 7,50E-03 | 1,89 | 1131 5 | 312 | 1135 | 59 |

**Table S4. Criteria for defining divergence classes following (Landry et al., 2005).**

| Divergence class | Alleles - Binomial test | Parents - Binomial test | ASE versus Parents - Fisher test | Additional |
|---|---|---|---|---|
| cis | ** (A1≠A2) | ** (P1≠ P2) | NS (P1/P2 = A1/A2) | NA |
| trans | NS (A1= A2) | ** (P1 ≠ P2) | ** (P1/P2 ≠ A1/A2) | NA |
| cis + trans | ** (A1≠A2) | ** (P1≠ P2) | ** (P1/P2 ≠ A1/A2) | log2(P1/P2)/log2(A1/A2) > 1 |
| cis - trans | ** (A1≠A2) | ** (P1≠ P2) | ** (P1/P2 ≠ A1/A2) | log2(P1/P2)/log2(A1/A2) < 1 |
| compensatory | ** (A1≠A2) | NS (P1 = P2) | ** (P1/P2 ≠ A1/A2) | NA |
| conserved | NS (A1 = A2) | NS (P1 = P2) | NS (P1/P2 = A1/A2) | NA |

A1 - allele 1, A2 - allele 2, P1 - parent 1, P2 - parent 2, ** - statistically significant with FDR 10%, NS - non-significant with FDR 10%

| Table S5. DNA sequencing yield. | | | |
|---|---|---|---|
| **Sample ID** | **Sampling location** | **Ecotype** | **Mapped reads *** |
| FC14 | Little Campbell River | Freshwater | 17438204 |
| FC13 | Little Campbell River | Freshwater | 23171229 |
| FC12 | Little Campbell River | Freshwater | 25146843 |
| FC16 | Little Campbell River | Freshwater | 30005091 |
| FC18 | Little Campbell River | Freshwater | 38399784 |
| FC15 | Little Campbell River | Freshwater | 15152985 |
| c363_P_FC08_F | Little Campbell River | Freshwater (parent) | 58346540 |
| LITC_DWN_4 | Little Campbell River | Marine | 28238410 |
| LITC_DWN_5_F | Little Campbell River | Marine | 37997779 |
| LITC_DWN_6_F | Little Campbell River | Marine | 46675891 |
| LITC_DWN_7_F | Little Campbell River | Marine | 27626027 |
| LITC_DWN_8_F | Little Campbell River | Marine | 30613683 |
| LITC_DWN_9_F | Little Campbell River | Marine | 25885479 |
| c363_P_FC18_M | Little Campbell River | Marine (parent) | 36326200 |
| Tyne8_27 | River Tyne | Freshwater | 57821343 |
| Tyne8_4 | River Tyne | Freshwater | 75419343 |
| Tyne8_7 | River Tyne | Freshwater | 80784624 |
| Tank422_4 | River Tyne | Freshwater | 120054557 |
| Tyne8_2 | River Tyne | Freshwater | 89618061 |
| Tyne8_1 | River Tyne | Freshwater | 58244138 |
| c172_P_533_M | River Tyne | Freshwater (parent) | 56617456 |
| Tyne2_16\|2015 | River Tyne | Marine | 89914271 |
| Tyne2_18 | River Tyne | Marine | 28775921 |
| Tyne2_20 | River Tyne | Marine | 55538908 |
| Tyne2_12 | River Tyne | Marine | 15830364 |
| Tyne2_16\|2014 | River Tyne | Marine | 24148885 |
| Tyne2_14 | River Tyne | Marine | 108126294 |
| c172_P_532_F | River Tyne | Marine (parent) | 66347185 |
| c214_P_512_M | River Shiel | Freshwater (parent) | 57188122 |
| c214_P_524_F | River Shiel | Marine (parent) | 61037784 |
| c212_P_551_M | River Forss | Freshwater (parent) | 53922480 |
| c212_P_454_F | River Forss | Marine (parent) | 27969046 |
| * samtools view -F 0x4 *.bam \| cut -f 1 \| sort \| uniq \| wc -l | | | |
